# Optimal Neuromuscular Performance Requires Motor Neuron Phosphagen Kinases

**DOI:** 10.1101/2025.03.18.643998

**Authors:** Karlis A. Justs, Danielle V. Latner (nee Riboul), Carlos D. Oliva, Yosuf Arab, Gabriel G. Bonassi, Olena Mahneva, Sarah Crill, Sergio Sempertegui, Paul A. Kirchman, Yaouen Fily, Gregory T. Macleod

## Abstract

Phosphagen systems are crucial for muscle bioenergetics - rapidly regenerating ATP to support the high metabolic demands of intense musculoskeletal activity. However, their roles in motor neurons that drive muscle contraction have received little attention. Here, we knocked down expression of the primary phosphagen kinase [Arginine Kinase 1; ArgK1] in *Drosophila* larval motor neurons and assessed the impact on presynaptic energy metabolism and neurotransmission *in situ*. Fluorescent metabolic probes showed a deficit in presynaptic energy metabolism and some glycolytic compensation. Glycolytic compensation was revealed through a faster elevation in lactate at high firing frequencies, and the accumulation of pyruvate subsequent to firing. Our performance assays included two tests of endurance: enforced cycles of presynaptic calcium pumping, and, separately, enforced body-wall contractions for extended periods. Neither test of endurance revealed deficits when ArgK1 was knocked down. The only performance deficits were detected at firing frequencies that approached, or exceeded, twice the firing frequencies recorded during fictive locomotion, where both electrophysiology and SynaptopHluorin imaging showed an inability to sustain neurotransmitter release. Our computational modeling of presynaptic bioenergetics indicates that the phosphagen system’s contribution to motor neuron performance is likely through the removal of ADP in microdomains close to sites of ATP hydrolysis, rather than the provision of a deeper reservoir of ATP. Taken together, these data demonstrate that, as in muscle fibers, motor neurons rely on phosphagen systems during activity that imposes intense energetic demands.

## Introduction

Phosphagen kinases play an integral role in the bioenergetics of skeletal and cardiac muscle where they catalyze the conversion of phosphagens to ATP, and back again (Wallimann et al., 1992). Phosphagen systems maintain deep reservoirs of phosphagens, such as phosphoarginine in fruit flies and phosphocreatine in mammals, and these reservoirs make ATP available at times of intense metabolic load acting as “temporal” buffers between energy supply and demand (Newsholme et al., 1978). ATP, and ultimately phosphagens, are subsequently regenerated by glycolysis and oxidative phosphorylation at a more leisurely pace (Hird, 1986). Phosphagen systems also play “spatial” buffering roles which ameliorate limitations caused by the relatively slow rates of ATP and ADP diffusion in the cytosol (Meyer et al., 1984). Benefits accrue through a close physical association between phosphagen kinases and both the sites of ATP production and the sites of ATP consumption. The colocalization of phosphagen kinases and the machinery of glycolysis (Kraft et al., 2000; Maughan et al., 2005) and oxidative phosphorylation [mitochondria; (Rojo et al., 1991; Wyss et al., 1992)] improves the efficiency of ATP synthesis as the kinases remove ATP as it is synthesized and regenerate ADP - the substrate for further ATP synthesis (Schlattner et al., 2016; Wallimann et al., 2011). Similarly, the colocalization of phosphagen kinases with pumps and other ATPases (Schlattner et al., 2016) provides benefits as the rapid removal of ADP and regeneration of ATP maintains the free energy available from ATP hydrolysis (Ellington, 1989; Iyengar, 1984; Wallimann et al., 1992). This phenomenon is especially evident in studies demonstrating that phosphagens can provide a more effective energy substrate than ATP itself (Korge et al., 1993; Minajeva et al., 1996; Xu et al., 1996). Strategic colocalization with phosphagen kinases dictates which ATPases get privileged access to ATP from the phosphagen reservoir (Linton et al., 2010; Meyer et al., 1984).

Phosphagen systems play an important role in nervous tissue, including central neurons (Andres et al., 2008; Lipton & Whittingham, 1982), and the removal of different creatine kinase (CK) isoforms leads to various behavioral and spatial learning deficits in mice (Jost et al., 2002; Streijger et al., 2004; Streijger et al., 2005). However, their role in neuronal bioenergetics and synaptic performance *per se* remains unclear. Previously, we established that mitochondrial-associated phosphagen kinases can be found in *Drosophila* motor neurons (MNs) and the lower MNs of mice (Justs et al., 2023), as reported for humans (Lowe et al., 2013). Furthermore, they are found in the MN axon terminals (Justs et al., 2023). In mammals, phosphagen kinases are represented by 5 CK genes, the products of which form isoenzymes that either reside in the cytosol or target to mitochondria (Bertin et al., 2007; Wallimann et al., 2011). Knock-out (KO) of mitochondrially-targeted isoenzymes show little impact on muscle performance phenotype (Steeghs, Heerschap, et al., 1997; Steeghs et al., 1998), while KO of cytosolic isoenzymes result in a nuanced phenotype where tetanus force is diminished but initial twitch force and endurance are relatively unaffected (Steeghs, Benders, et al., 1997; van Deursen et al., 1993). In *Drosophila*, phosphagen kinases are represented by 3 arginine kinase (ArgK) genes (CG4929, CG5144 and CG4546), with only one (CG4929; ArgK1) showing significant expression in the larval nervous system (flybase.org). While we previously demonstrated that knock-down (KD) of *Drosophila* ArgK1 compromises presynaptic energy metabolism (Justs et al., 2023), we did not investigate the extent to which neuromuscular performance relies upon an intact phosphagen system.

Here, we developed a number of imaging techniques to determine the impact of ArgK1 KD on presynaptic energy metabolism, the ability of motor nerve terminals to pump calcium ions (Ca^2+^) during burst firing, and the ability of larvae to sustain body-wall contractions. Using a new suite of fluorescent probes (ATeam^YEMK^, LiLac and Pyronic) we demonstrated the impact of ArgK1 KD on ATP, lactate and pyruvate levels and found the data to be consistent with previous observations (Justs et al., 2023). Our electrophysiological assays only revealed deficits in neurotransmission during high frequency nerve stimulation. To address the difficulty of studying neurotransmission at neuromuscular junctions (NMJs) under physiological conditions - such as repeated high-frequency bursts, where muscle contractions displace electrodes - we developed two MN performance assays to simulate *in vivo* firing. Our findings suggest that the ArgK1-enabled phosphagen system in MNs has minimal effect on endurance but may influence the power exerted at the start of body wall contractions.

## Results

### A small portion of ArgK1 localizes to presynaptic mitochondria via an N-terminal signal sequence

In *Drosophila*, ArgK1 is alternatively spliced, and transposon-mediated insertion of GFP DNA into an intron of ArgK1 via MiMIC cassette (stock #51522, BDSC; Fig.1A) revealed that at least 1 of the 7 reported isoforms target to mitochondria [Fig.1B-C; Justs et al. (2023)]. To test whether one or both N-terminal exons are required for mitochondrial targeting, we made two transgenic animals:-one in which oxBFP DNA was C-terminally appended to cDNA coding the first exon, and one in which oxBFP was flanked by cDNA of the first two exons (Fig.1D). Both UAS constructs targeted the fluorophores to mitochondria, demonstrating that the first exon was sufficient (Fig.1D-F). Unexpectedly, the first exon in the absence of the second resulted in mitochondria that appeared to “round up” (Fig.1E), indicating a dominant-negative effect of the exogenous first exon peptide (MFALWYLTFAVDEIRK). This analysis indicated that 2 of the 7 ArgK1 splice isoforms (those including the first exon) localize to mitochondria.

**Figure 1.**
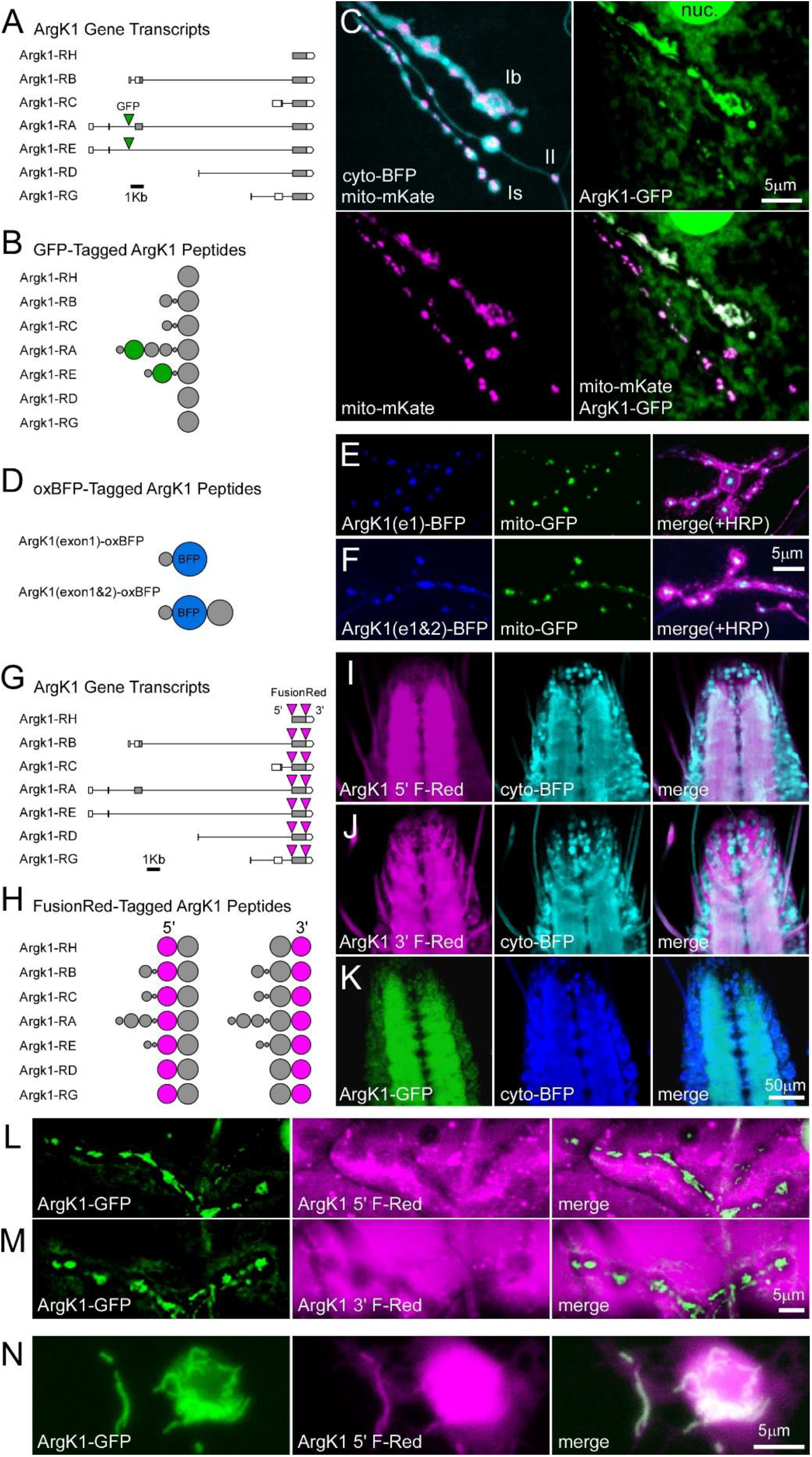
ArgK1 is targeted to presynaptic mitochondria by an N-terminal sequence. **A**. Predicted splice isoforms of ArgK1 (FlyBase) showing intronic sites of transposon-mediated GFP DNA insertion via MiMIC cassette. **B**. Representation of the proteins generated from ArgK1 splice isoforms, where the volume of each ‘sphere’ represents the relative length of peptide corresponding to each exon. **C**. An image of a short confocal stack projection through live MN (MN) terminals of MN13-Ib and MNSNb/d-Is and MNSNb/d-II on muscle fiber #13, demonstrating the relative localization of ArgK1-GFP (1 copy) and neuronal expression of mitochondrial-targeted mKate [UAS-mKate; Justs et al. (2023)] and cytosolic TagBFP. **D**. Representation of the proteins generated from two different transgenic constructs; ArgK1(exon1)-oxBFP where oxBFP DNA was C-terminally appended to DNA coding the first ArgK1 exon, and ArgK1(exon1)-oxBFP where oxBFP was flanked by DNA of the first two exons of ArgK1. **E**. An image (single confocal plane) of live MN terminals showing ArgK1(exon1)-oxBFP localization relative to neuronal expression of mitochondrial-targeted GFP (mito-GFP) and a Cy3-conjugated HRP antibody (HRP-Cy3). **F**. An image (single confocal plane) of live MN terminals showing ArgK1(exon1&2)-oxBFP localization relative to mito-GFP and an HRP-Cy3 antibody. **G**. Predicted splice isoforms of ArgK1 showing Fusion Red DNA insertion sites (5’ or 3’) in two different CRISPR/Cas9-edited fly lines. One fly line contains FusionRed DNA inserted at the N-terminus of the last exon (ArgK1 5’ Fusion Red), while the other has FusionRed DNA inserted at the C-terminus of the last exon (ArgK1 3’ Fusion Red). **H**. Representation of the proteins generated in each of the two fly lines with endogenously-tagged ArgK1. **I**. An image (single confocal plane) of the live ventral ganglion (VG) showing the neuropil localization of ArgK1 5’ Fusion Red relative to pan-neuronal expression of cytosolic TagBFP. **J**. An image of the VG showing ArgK1 3’ Fusion Red relative to cytosolic TagBFP. **K**. An image of the VG showing ArgK1-GFP expression relative to cytosolic TagBFP. **L**. An image of a short confocal stack projection through live terminals of MN13-Ib and MNSNb/d-Is, showing a strong presence of ArgK1 5’ F-Red (1 copy) in muscle fiber #13, but little sign of Fusion Red in the mitochondria of MN terminals revealed by the presence ArgK1-GFP (1 copy). **M**. As in L, but in the ArgK1 3’ F-Red line. **N**. ArgK1 5’ F-Red colocalized with ArgK1-GFP revealed mitochondria in live MNs cultured from the ventral ganglion of larvae bearing one copy of the ArgK1 5’ F-Red edited allele and one copy of the ArgK1-GFP allele (as in L). nSyb-GAL4 was used to drive expression of transgenes in panels C-K.

To reveal the expression pattern of all ArgK1 splice isoforms we used the CRISPR/Cas9 gene editing technique to insert FusionRed DNA at either the 5’ or the 3’ end of the catalytic portion of ArgK1 (Fig.1G-H). FusionRed revealed an endogenous ArgK1 expression pattern that filled the neuropil of the larval ventral ganglion (Fig.1I-J), similar to the expression pattern of ArgK1-GFP (Fig.1K). However, the expression pattern was difficult to interpret when looking at the NMJ as FusionRed in the muscle overwhelmed any FusionRed that might have been detected in MN terminals (Fig.1L-M). This ambiguity was settled by examining primary cell cultures, where FusionRed was clearly colocalized with MN mitochondria (Fig.1N).

### ArgK1 KD leads to deficits in presynaptic energy metabolism

Using a fluorescent reporter of the cytosolic ATP/ADP ratio (PercevalHR) we previously demonstrated that ATP/ADP levels fall further in ArgK1 KD terminals than in controls under intense metabolic load (Justs et al., 2023). However, as PercevalHR is pH sensitive, and these MN terminals acidify under load (Rossano et al., 2013), we adopted the use of ATeam1.03^YEMK^ (Imamura et al., 2009), a reporter of ATP levels less susceptible to pH change near neutral, with an intracellular dissociation constant (K_D_) of ~2.7 mM at room temperature *in situ* (Lerchundi et al., 2020). To impose a robust metabolic load, hemisegment nerves containing MN axons were stimulated for 10 seconds at a frequency nearly double their native firing frequency (80Hz). The presence of physiological levels of extracellular Ca^2+^ obligates physiological levels of Ca^2+^ pumping and neurotransmitter release, but muscle contraction can be stifled by adding glutamate (7mM) to desensitize postsynaptic glutamate receptors (Macleod et al., 2004).

ATeam1.03^YEMK^ revealed a rapid reduction in ATP levels and ATP levels appeared to fall further in ArgK1 KD terminals (Fig.2A-E), but the trend was not found to be significant at its maximum extent (P = 0.052). Importantly, the ATP levels measured here are expected to fall less sharply than the ATP/ADP ratio reported previously (Justs et al., 2023).

**Figure 2.**
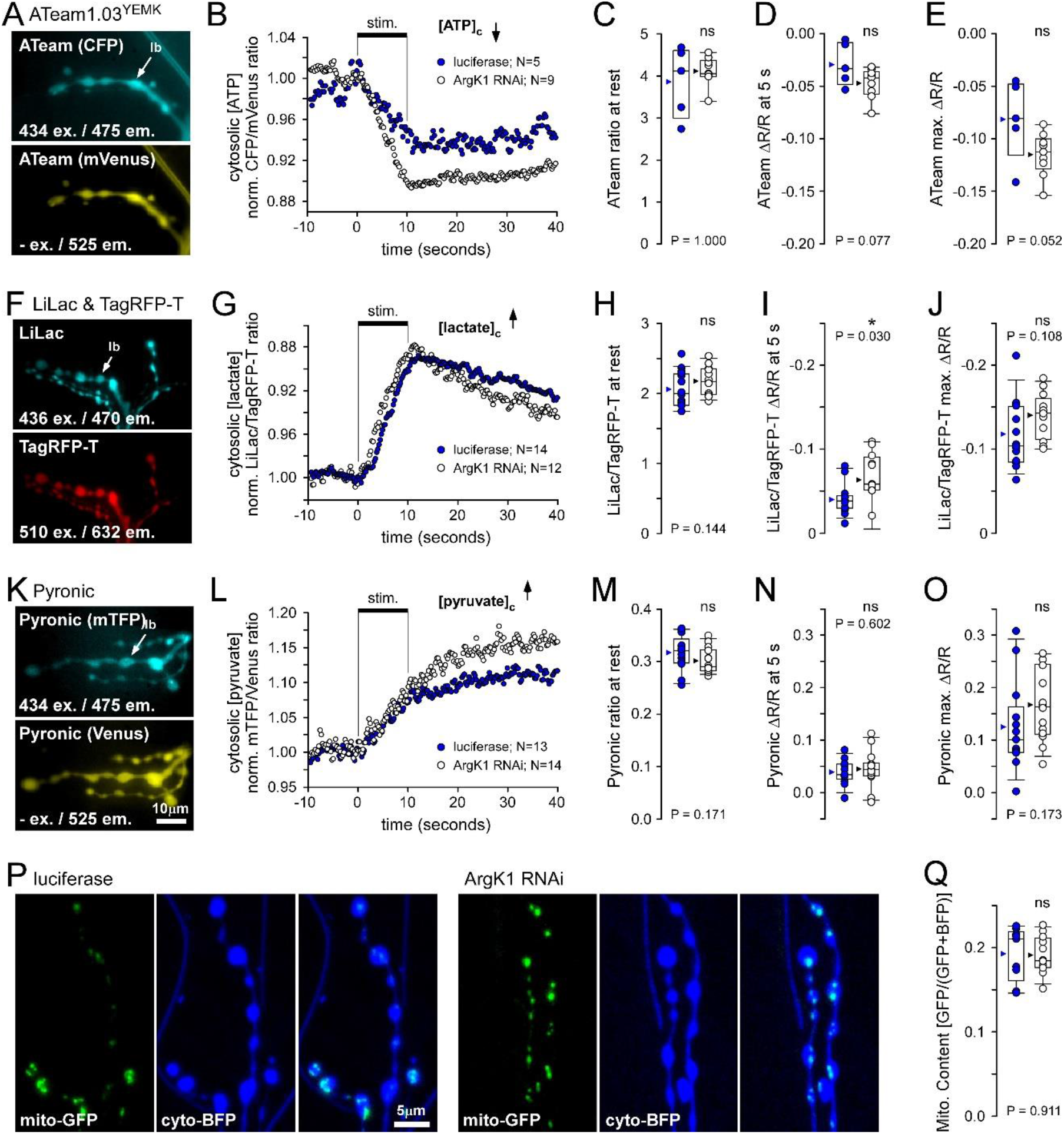
Presynaptic Metabolic Imaging and Mitochondrial Content after ArgK1 KD. **A**. Images of motor nerve (MN) terminals on muscle fiber #13 expressing ATP reporter ATeam1.03^YEMK^ captured prior to stimulation using excitation and emission wavelengths as shown (mVenus is excited by CFP emission). Pan-neuronal nSyb-GAL4 was used to drive UAS-ATeam1.03^YEMK^ expression along with either UAS-ArgK1 dsDNA, to knock down ArgK1, or UAS-luciferase as a control. **B**. A plot showing the normalized average ATeam1.03^YEMK^ fluorescence ratio measured at MN13-Ib terminals while stimulated at 80Hz for 10 seconds, as shown. A ratio decrease is interpreted as a decrease in the cytosolic ATP concentration ([ATP]_c_). The numbers of independent preparations are indicated. **C**. Box plots of ATeam1.03^YEMK^ ratio prior to nerve stimulation (at rest) for each preparation. **D**. Change in ATeam1.03^YEMK^ ratio (ΔR/R_rest_) 5 seconds after start of stimulus train. **E**. Greatest change in the ATeam1.03^YEMK^ ratio (max ΔR/R_rest_); minimum detected between 8 and 12 seconds. **F**. Images of MN terminals expressing lactate reporter LiLac::TagRFP-T captured prior to nerve stimulation. **G**. The normalized average LiLac to TagRPF-T ratio shown responding to nerve stimulation. Scale is inverted and lower numbers are interpreted as higher concentrations of lactate ([lactate]_c_). **H**. LiLac to TagRFP-T ratio at rest. **I**. Change in LiLac to TagRFP-T ratio (ΔR/R_rest_) 5 seconds after start of stimulus train. **J**. Greatest change in the LiLac to TagRFP-T ratio (max ΔR/R_rest_); maximum detected between 8 and 12 seconds. **K**. Images of MN terminals expressing pyruvate reporter Pyronic, captured prior to stimulation. Venus is excited by mTFP emission. Scale bar applies to images in A, F and K. **L**. The normalized average Pyronic ratio shown responding to nerve stimulation. An increase in the ratio is interpreted as an increase in pyruvate concentration ([pyruvate]_c_). **M**. Pyronic ratio at rest. **N**. Change in Pyronic ratio (ΔR/R_rest_) 5 seconds after start of stimulus train. **O**. Greatest change in the Pyronic ratio (max ΔR/R_rest_); measured at 40 seconds. nSyb-GAL4 was used to drive pan-neuronal expression of all metabolic reporter transgenes. All imaging metabolic imaging was performed on the terminal of MN13-Ib in segment #4. Average traces shown. A three-point moving average was used on ATeam1.03^YEMK^ and Pyronic traces. ATeam1.03^YEMK^ fluorescence ratio (but not LiLac or Pyronic) corrected for bleaching as described in the Methods. **P**. Images from short confocal stack projection through live terminals on muscle fiber #6. OK6-GAL4 was used to drive UAS-mito-GFP and UAS-cyto-BFP expression in MNs, along with either UAS-ArgK1 dsDNA to knock down ArgK1 (N=12) or UAS-luciferase as a control (N=9). **Q**. Box plots of mitochondrial content of MN6/7-Ib terminals. Mitochondrial content was quantified as the amount of mito-GFP fluorescence divided by the sum of mito-GFP and cyto-BFP fluorescence. Units are arbitrary. Experiments were discarded as outliers if the initial ratio of ΔR/R was beyond 3 x median absolute deviations of the median in C-E, H-J and M-O. Box plots show values from all preparations; mean (arrowhead), median (line), 25-75 percentiles box and 10-90% whiskers. Mann-Whitney Rank Sum test in C and H; unpaired Student’s t-tests used on all other two sample comparisons.

Previously, we found that lactate levels increase at a faster rate in ArgK1 KD terminals than in controls under an intense metabolic load, and we speculated that this is a result of either a compensatory increase in glycolysis or a reduced mitochondrial ability to metabolize pyruvate, leading to a faster rate of pyruvate conversion to lactate (Justs et al., 2023). Here, we used an improved fluorescent cytosolic reporter of lactate [LiLac (Koveal et al., 2020)] to re-examine changes in lactate levels (Fig.2F-J) and we used cytosolic pyruvate reporter Pyronic (Gonzalez-Gutierrez et al., 2020; San Martin et al., 2014) to make complementary measurements of presynaptic pyruvate levels (Fig.2K-O). LiLac measurements, like previous Laconic measurements, revealed faster rates of lactate accumulation in ArgK1 KD terminals than in controls (P = 0.030) (Fig.2I), and while Pyronic suggested that pyruvate levels exceeded control levels after nerve stimulation, the trend did not reach statistical significance (P = 0.173) (Fig.2O). An increase in pyruvate would be consistent with mitochondria being unable to keep pace with the rate of pyruvate production in ArgK1 KD terminals, but ultimately, the data did not provide the statistical basis for rejecting the alternate hypothesis that there was a compensatory increase in glycolysis and pyruvate production.

In mammals, knock-down of various isoforms of CK leads to an increase in the glycogen content and glycolytic capacity of skeletal muscle fibers, as well as an increase in mitochondria content and the expression and activity of respiration chain proteins (de Groof et al., 2001; Dzeja et al., 2011; Kaasik et al., 2003; Steeghs, Benders, et al., 1997; Veksler et al., 1995).

Here, we examined MN terminals for an increase in mitochondrial content in response to ArgK1 KD. MN6/7-Ib terminals were examined as they have a high energy demand and mitochondrial volume density (6.29%; Justs et al. (2022)). Fluorescence microscopy, although poorly suited for making quantitative estimates of mitochondrial content, is well suited for estimating *changes* in content. Our measure of content was calculated as the mitochondrial fluorescence intensity divided by the sum of mitochondrial and cytosolic fluorescence intensity (different fluorophores) (Fig.2P-Q). ArgK1 KD did not result in a significant change in the mitochondrial content reported by this ratio, whether fluorophore expression was driven by OK6-GAL4 (P=0.911; Fig.2Q) or a stronger pan-neuronal driver (nSyb-GAL4; P=0.577; data not shown).

### Deficits in neurotransmission are revealed by burst firing after ArgK1 KD

As an initial test of the impact of knocking down ArgK1 on neurotransmitter release, we used single muscle fiber voltage clamp and current clamp techniques to analyze neurotransmitter release from two MNs that synapse on muscle fiber #6 (Fig.3A). During fictive locomotion MN6/7-Ib fires at 21.4Hz, while MNSNb/d-Is fires at 7.8Hz (Justs et al., 2022). We chose a supramaximal stimulus frequency for the two MNs (60Hz; Fig.3B), one that might reasonably test the capacity of the MNs to sustain neurotransmitter release if energy metabolism is impaired, and we maintained stimulation for approximately half the period expected for a contraction cycle during unrestrained locomotion (Fig.3C). The amplitude of the *compound* excitatory junctional current (EJC) was not significantly different at the start of the stimulus train (Fig.3D; P=0.537), but neurotransmission depressed more rapidly when ArgK1 was knocked down (Fig.3E; P<0.001).

**Figure 3.**
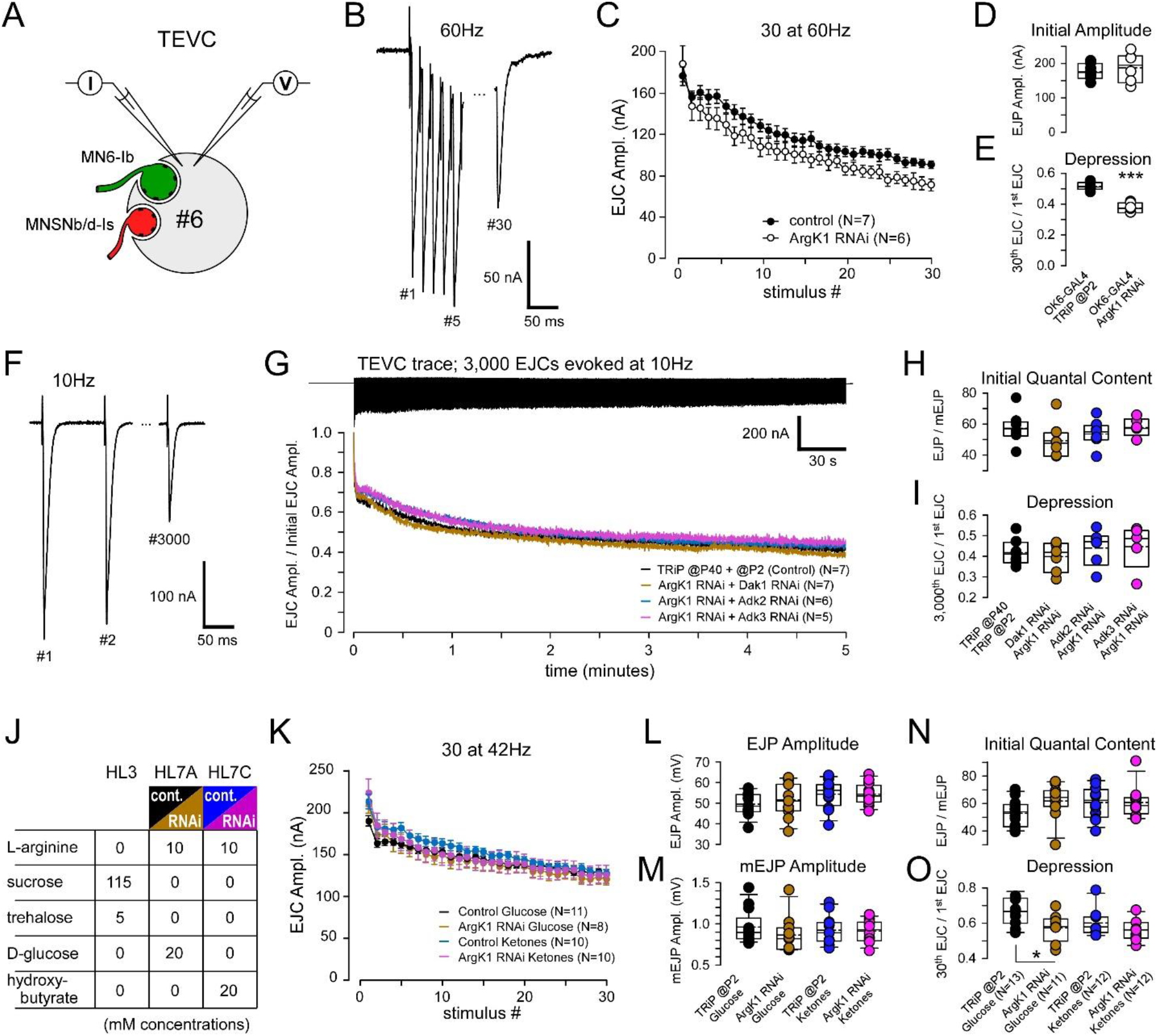
Electrophysiological analysis of the impact of ArgK1 KD on neurotransmission. **A**. A diagram of a transverse section through muscle fiber #6, innervated by MNs MN6/7-Ib and MNSNb/d-Is, and subject to a two-electrode voltage clamp (TEVC). **B**. Example trace showing compound excitatory junctional currents (EJCs). The hemisegment nerve was stimulated at a supramaximal rate (60Hz), well above their individual endogenous firing rates. **C**. Plots of average compound EJC amplitude when the nerve was stimulated for 500ms. ArgK1 expression was knocked down in MNs by OK6-GAL4 driven expression of ArgK1 dsRNA (BL#41697), with OK6-GAL4 driving P{CaryP}attP2 (BL#36303) as a control. Recordings in B-E were conducted in HL3 saline; 3 mM [Ca^2+^] and 20 mM [Mg^2+^]. Experiments were discarded as outliers if the initial amplitude or depression was beyond 3 x median absolute deviations of the median. **D**. Plots of compound EJC amplitude in response to the first stimulus in the train in C. Student’s t-test (not sig., P=0.537). **E**. Depression ratio box plots (ratio of amplitudes between 30^th^ and 1^st^ EJC). Student’s t-test (***, P<0.001). **F**. Example trace showing compound EJCs in muscle #6 while stimulating the hemisegment nerve at 10Hz. **G**. Top: Example trace showing compound EJCs in muscle #6 while stimulating the nerve at 10Hz for 5 minutes (3,000 impulses). Bottom: Plots of average EJC amplitude normalized to the amplitude of the first EJC in each train. In each case ArgK1 expression was knocked down by nSyb-GAL4 driven expression of dsRNA in the MNs, combined with dsRNA for knockdown of Dak1, Adk2 or Adk3. As a control, nSyb-GAL4 drove both P{CaryP}attP40 (BL#36304) and P{CaryP}attP2 (BL#36303). Recordings in F-I were conducted in HL6 saline (2 mM [Ca^2+^] and 15 mM [Mg^2+^]) to best maintain the preparation during prolonged stimulation. **H**. Single electrode current-clamp recordings determined the quantal content (QC) of release evoked from both terminals, prior to the insertion of a second electrode to implement a two-electrode voltage clamp (TEVC). Box plots of the average QC, calculated by dividing the average corrected EJP amplitude (≥10 EJPs evoked at 0.2Hz) by the average amplitude of ≥30 miniature EJPs (mEJPs). **I**. Depression ratio box plots (ratio of amplitudes between 3,000^th^ and 1^st^ EJC). No significant differences were found by one-way ANOVA applied in H (P=0.336) or I (P=0.725). **J**. A table highlighting differences in salines used to test the susceptibility of neurotransmission to metabolic substrates that reduce the opportunity for glycolytic compensation. L-arginine was increased to accommodate the possibility of a greater ArgK1 substrate requirement for ATP buffering. **K**. Average compound EJC amplitude when the nerve was stimulated at 42Hz for 700ms (30 impulses). The stimulus duration reflects a combination of the typical firing periods for MN6/7-Ib (830ms) and MNSNb/d-Is (210ms) during a 1-second peristaltic contraction (Justs et al., 2022). ArgK1 expression was knocked down by nSyb-GAL4 driven expression of ArgK1 dsRNA, with P{CaryP}attP2 (BL#36303) driven as a control. Recordings were conducted in one of the salines listed in J; each a derivative of HL3 that also included (in mM) 5 KCl, 80 NaCl, 16 NaHCO_3_, 2 CaCl_2_ and 15 MgCl_2_. **L**. Box plots showing the average amplitude of compound EJPs evoked at 0.2Hz. A one-way ANOVA detected no significant differences (P=0.201). **M**. Box plots of the average amplitude of mEJPs collected during the same recordings used to collect EJPs (N). No significant differences were found by Kruskal-Wallis one-way ANOVA on ranks (P=0.551). **N**. Box plots of the average QC calculated as in H, using the corrected EJP and mEJP data shown in L and M (current clamp). A one-way ANOVA detected no significant differences (P=0.185). **O**. Depression ratio box plots (ratio of 30^th^ to the 1^st^ EJC). An asterisk indicates significantly greater depression when ArgK1 was knocked down (P < 0.014); one-way ANOVA with Holm-Sidak post hoc tests. Box plots show mean (dotted line) and median, with 25-75% boxes and 5-95% whiskers. OK6-GAL4 was used to expression of transgenes in panels B-E, while nSyb-GAL4 was used in F-O.

We then sought to determine whether the phosphagen system plays a role in sustaining release over longer time periods. Physiological assays testing a reliance on energy metabolism require physiologically relevant loads and these can only be induced using high levels of extracellular Ca^2+^. However, high Ca^2+^ levels evoke massive muscle contraction at high stimulus frequencies, and recordings cannot be maintained beyond fractions of a second. We therefore chose a lower stimulus frequency (10Hz; Fig.3F), one that would allow us to maintain a two-electrode voltage clamp (TEVC) over several minutes (Fig.3G). To guard against the possibility that other ATP regeneration mechanisms compensate when ArgK1 is knocked down, we knocked down adenylate kinases (AKs) in parallel. In mammals, inhibition of CKs has been shown to increases the phosphotransfer flux through AKs (Dzeja et al., 1996). There are three AK isozymes in mammals (Dzeja & Terzic, 2009), and three have been identified in *Drosophila* (Dak1, Adk2 and Adk3; flybase.org) for which RNAi lines are available. Our analysis of quantal content (QC), gleaned from current-clamp recordings at 0.2Hz prior to TEVC recordings, showed that the initial evoked release was similar between conditions knocking down Argk1 and alternate AKs (Fig.3H; P=0.336). We were unable to detect any deficit in neurotransmission over a sustained period of 10Hz stimulation (Fig.3I; P=0.725).

As we observed that glycolytic activity increased under load when we knocked down ArgK1 (Fig.2), we sought to determine the full impact on neurotransmission under conditions where glycolytic activity is unable to compensate. To minimize the potential for glycolytic compensation we replaced carbohydrates (glucose) in the saline with fatty acids (ketone bodies: hydroxybutyrate) (Fig.3J) which are available for oxidative phosphorylation through fatty acid metabolism, but not glycolysis. A stimulus frequency of 42Hz was adopted as the supramaximal frequency (Fig.3K) as the experimental failure rate at 60Hz (Fig.3C) proved to be too high to collect enough data for each of the four treatments. We found no difference in the amplitude of evoked EJPs, mEJPs or the calculated QC prior to implementation of a TEVC or train stimulation (Fig.3L, P=0.201; 3M, P=0.551; 3N, P=0.185). Furthermore, although we still observed more rapid depression in neurotransmission when ArgK1 was knocked down (Fig.3O; P=0.014), we saw no greater depression in the absence of glucose that limited glycolytic compensation.

### Ca^2+^ pumping during burst firing is not impaired by ArgK1 KD

Ultimately, the metabolic loads we can impose while monitoring neurotransmission are moderate at best, due to the muscle contraction that occurs when nerves are stimulated at high frequency in the presence of physiological levels of extracellular Ca^2+^. Glutamate, added to the bath to stifle muscle contraction during metabolic imaging, cannot be used, as this extinguishes the electrophysiological signs of neurotransmitter release. Using Ca^2+^ imaging techniques we tested the capacity of ArgK1 KD terminals to pump Ca^2+^ and function under conditions of a metabolic load beyond that we could impose during electrophysiological recordings. MNs, expressing the genetically-encoded ratiometric Ca^2+^ indicator mScar8f (Li et al., 2021), were stimulated at 50Hz for 2 seconds at 4 second intervals, for a total of 4 minutes (Fig.4A-C).

Importantly, although it was only Ca^2+^ cycling that was assessed, the same stimulus trains initiate action potentials and trigger neurotransmitter release imposing a substantial metabolic burden as it obligates the recycling and refilling of synaptic vesicles (SVs) in addition to resetting Na^+^, K^+^ and Ca^2+^ gradients across the plasma membrane. Our readout of the terminals ability to function was assessed through its ability to remove Ca^2+^ in response to each stimulus cycle (Fig.4C). Our expectation was that, beyond a certain metabolic load in ArgK1 KD terminals, ATP levels would fall below a particular threshold and Ca^2+^ levels would become dysregulated.

Despite a robust metabolic load, Ca^2+^ cycling appeared to be as efficient in ArgK1 KD terminals as in control terminals (Fig.4D-F).

To confirm that this assay has the power to detect a deficit in energy metabolism we repeated the assay in the presence of extracellular Sr^2+^ rather than Ca^2+^ (Fig.4G-J). As at mammalian presynaptic terminals, Sr^2+^ permeates *Drosophila* VGCCs (Chouhan et al., 2012) where it triggers neurotransmitter release (Jan & Jan, 1976). Therefore, Sr^2+^ will support activity that imposes a substantial metabolic demand, but as Sr^2+^ is less effective at activating Krebs cycle enzymes (McCormack & Osbaldeston, 1990) oxidative phosphorylation and energy supply will be compromised. We observed greater depression of cycling in the presence of Sr^2+^ relative to Ca^2+^ (Fig.4J; P=0.010), indicating that the absence of a Ca^2+^ cycling deficit in ArgK1 KD terminals was due to the absence of a deficit, rather the absence of a capacity to detect a deficit.

### ArgK1 KD results in minimal impairment of musculoskeletal performance

While the cyclical Ca^2+^ pumping protocol demonstrated in Figure 4 imposed a metabolic load beyond levels we could impose during electrophysiology, the load was still only moderate in the context of what might be expected *in vivo*. During locomotion, the MN13-Ib MN likely fires at around 42.0Hz (Chouhan et al., 2012), with a duty cycle of 0.8 (Klose et al., 2005), resulting in 8,160 APs over a period of 4 minutes. However, we only imposed 6,060 impulses in the Ca^2+^ cycling protocol (50Hz x 2 sec x 240 / 4). Longer Ca^2+^ cycling protocols were problematic, leading to fluorophore bleaching, phototoxicity and difficulties in maintaining focus. We were also unable to impose higher firing rates, or more closely spaced cycles as it diminished our ability to quantify the cycles. Therefore, to challenge a MN’s capacity to maintain neurotransmission, not just Ca^2+^ cycling, over longer periods of time, we used opsins to excite MNs and we monitored larval contraction. Significantly, this *in vivo* assay tests MN performance while sidestepping the confound of motivation to perform.

**Figure 4.**
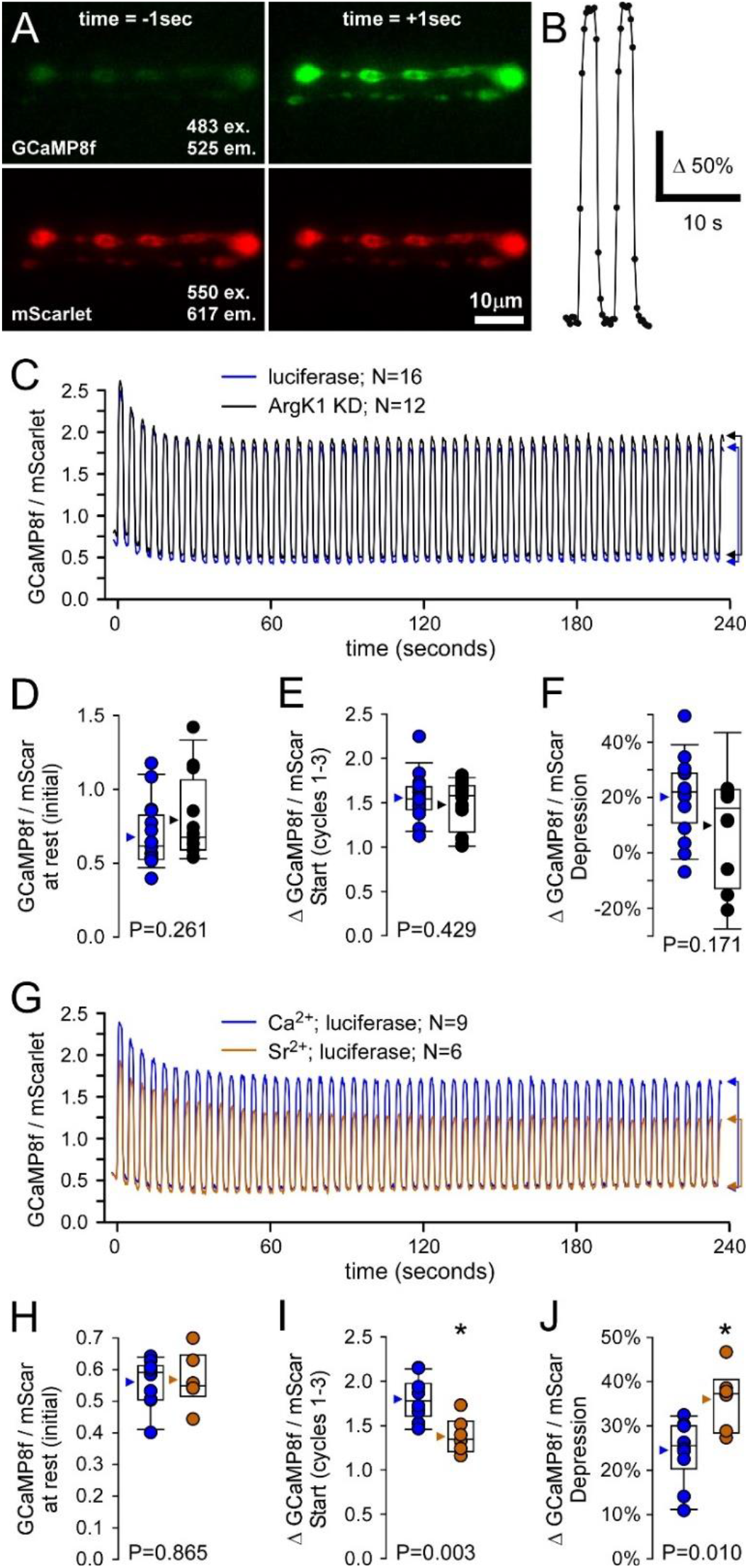
Imaging of presynaptic Ca^2+^ levels and Ca^2+^ pumping capacity after ArgK1 KD. **A**. Images of motor neuron (MN) terminals on muscle fiber #13 expressing the ratiometric Ca^2+^ indicator mScar8f (GCaMP8f fused to mScarlet and the SV protein synaptotagmin). Images were taken before and after 50Hz nerve stimulation, using the specified excitation and emission wavelengths. **B**. An example trace of the GCaMP8f to mScarlet fluorescence intensity ratio during two cycles of nerve stimulation, taken from a single trial (N=1) without signal averaging. Images pairs were collected every 250 ms to generate a single ratio measurement (point). An increase in the ratio indicates an increase in the cytosolic Ca^2+^ concentration ([Ca^2+^]_c_). **C**. Plots showing the average GCaMP8f to mScarlet ratio over 55 cycles from larvae in which ArgK1 has been knocked down [UAS-ArgK1 dsDNA; N=12] and control larvae in which the same MN driver is used to express luciferase [UAS-luciferase; (N=16)]. **D**. Plots of the ratio prior to nerve stimulation (at rest) from each preparation. **E**. Plots of the average amplitude of the ratio response over the first 3 cycles of the 55-cycle protocol. **F**. Plots of the change (depression) in the amplitude of the ratio between the first and last 3 cycles of the 55-cycle protocol. Data in A-F collected in HL6 with 2mM CaCl_2_, 15mM MgCl2 and 7mM L-glutamate. **G**. Plots of the ratio during cycles of nerve stimulation in the HL6 described above (N=9), and with 4mM SrCl_2_ and 2mM EGTA in place of 2mM CaCl_2_ (N=6). Control larvae expressing luciferase used for both Ca^2+^ and Sr^2+^ conditions. **H-J**. Plots as in D-F. Asterisks denote significant differences as determined by Student’s T-test. Box plots show mean (dotted line) and median, with 25-75% boxes and 5-95% whiskers. nSyb-GAL4 was used to drive expression of transgenes.

A mutated form of channelrhodopsin-2 [H134R-ChR2; (Nagel et al., 2005; Pulver et al., 2009)], which is activated by blue light (Fig.5A), was expressed using a MN-specific driver (OK371-GAL4). The larvae were illuminated for 2 seconds at 3 second intervals, causing immediate, robust, and consistent cycles of larval contraction (Fig.5B). Rest periods between cycles of illumination were brief (1 second) to allow recovery but without providing larvae the opportunity to locomote out of the field of view. Individual larvae were submitted to cyclical illumination for 20 minutes before data were analyzed and pooled (Fig.5C). During the development of this assay, we demonstrated that larvae with a reduced volume of mitochondria in their MN terminals (*dmiro* KD) were unable to sustain contractions, confirming the capacity of this assay to detect contraction deficits arising from impaired energy metabolism in MNs (Arab et al., 2025). After 20 minutes, ArgK1 KD larvae appeared to be cycling between states that were marginally more contracted than the control larvae (Fig.5E; Fig.5I, relaxed, P=0.197; Fig.5J, contracted, P=0.018), yet we were at a loss to explain how such a phenotype might arise. Ultimately, however, the extent of ArgK1 KD larval contraction was indistinguishable from control over the first 10 cycles (Fig.5D, F & K; P=0.930), and ArgK1 KD larval contraction was no more depressed than in control larvae after 20 minutes (Fig.5E, G & L; P=0.317).

**Figure 5.**
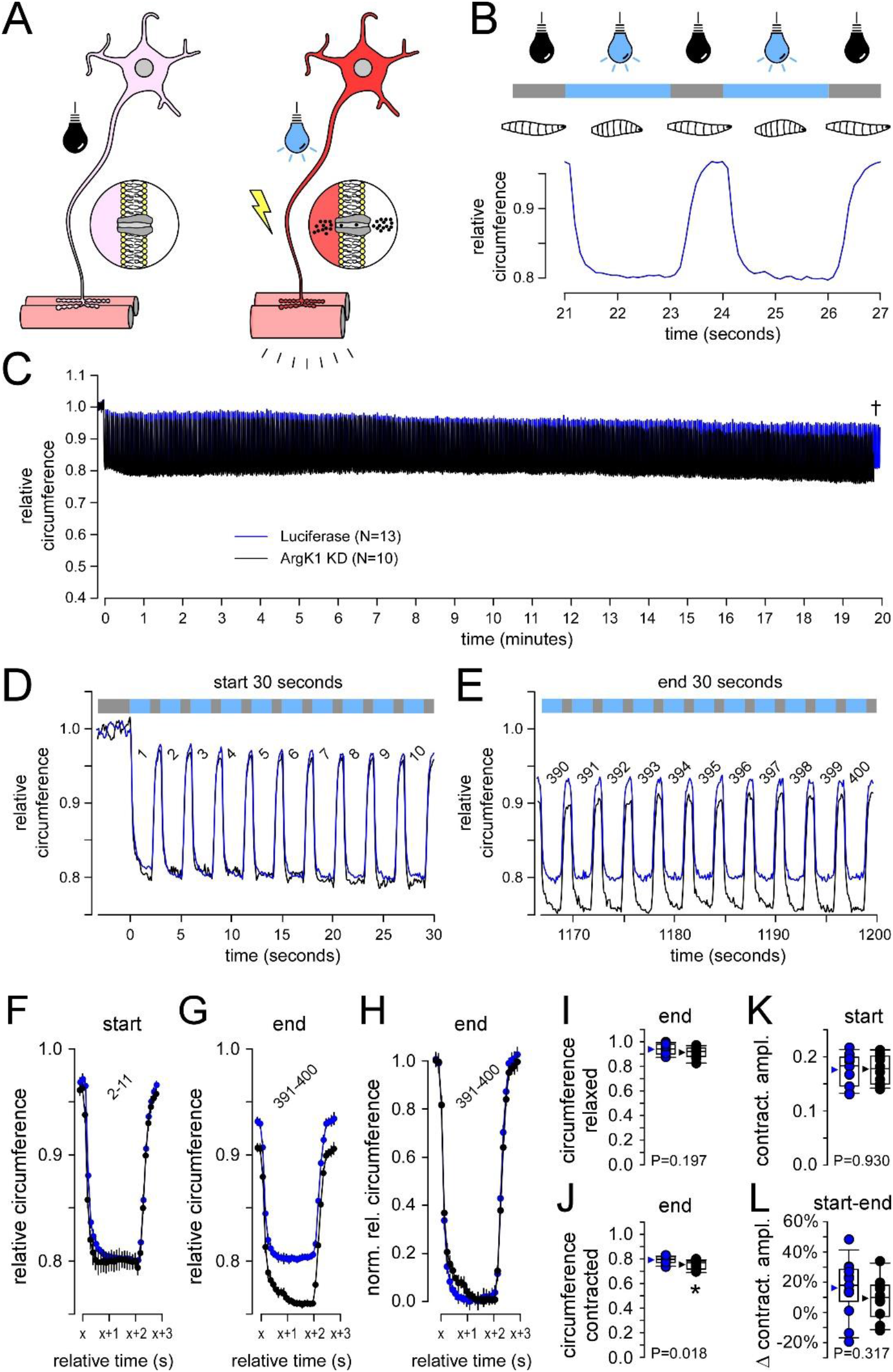
Analysis of musculoskeletal performance after ArgK1 KD. **A**. Channelrhodopsin-2 (ChR2) expressed in MNs (MNs) can be light activated to cause MN firing, neurotransmitter release and muscle contraction. **B**. Cyclical bouts of blue light illumination is effective at causing immediate and sustained larval contraction that ceases upon cessation of illumination. **C**. A normalized plot of cyclical larval contraction showing the average perimeter measurement of individual larvae in which ArgK1 KD has been knocked down [UAS-ArgK1 dsDNA; N=10; black trace], and control in which the same MN driver (OK371-GAL4) is used to express luciferase [UAS-luciferase; N=13; blue trace]. The dagger indicates that the final 10 seconds of the ArgK1 KD trace is omitted to allow a comparison between larvae. **D**. A plot of the detail of the first 10 cycles (numbered) of larval contraction taken from C. **E**. Detail of the last 10 cycles (numbered) of larval contraction taken from C. **F**. A plot of the average of contraction cycles 2-11 in C. Error bars show standard deviation in F- **G**. The average of the last 10 contraction cycles (391-400) in C. **H**. The plots in G normalized to the average of the first two data points in each trace. **I-J**. Box plots of the relative circumference of the larvae at maximum rest between cycles (I) and maximum contraction during cycles (J) over the last 10 contraction cycles. **K**. Box plots of the change in the normalized perimeter between rest and maximum contraction, for each larva, over contraction cycles 2-11. **L**. Box plots of change in the contraction amplitude over the last 10 contraction cycles compared to cycles 2-11. Values are shown from all preparations; mean (arrowhead), median (line), 25-75 percentiles box and 10-90% whiskers. Student’s T tests are used to test for significance in plots I-L. OK371-GAL4 (two copies) were used to drive expression of transgenes.

### Simulations of [ATP] during endurance assays show little depletion in the absence of a phosphagen system

The experiments above (Figs.3-5) proceeded under the assumption that a neuronal phosphagen system is essential for sustaining presynaptic activity. However, to the extent that we could quantify presynaptic performance while driving MNs at high frequencies, we detected little evidence of deficits. To better understand the bioenergetic capacity of these MNs we simulated ATP kinetics in a previously described computation model (Justs et al., 2023) (Fig.6). The model calculates total energy *demand* during presynaptic activity based on empirical estimates of neurotransmitter release and Ca^2+^ entry and based on theoretical estimates of Na^+^ entry [Fig.6A-C; Justs et al. (2022). It calculates ATP production according to the stimulatory influence of the energy state ([ATP]/([ADP][P_i_])]; Wilson (2017)) while limited by production maxima drawn from empirical estimates of oxidative phosphorylation from larval (0.079 mM/s/1%) and adult tissues (0.154 mM/s/1%) [Fig.6D; (Menail et al., 2022; Neville et al., 2018)]. The model is therefore able to simulate the levels of adenine nucleotides (ATP, ADP and AMP) and inorganic phosphate (P_i_) net of demand and production, and their rate of change in the presence of a phosphagen system or in its absence (Fig.6E-I; 6L-P).

The simulations indicate that ATP levels could be maintained if the MNs had an ATP production limit as high as 0.154 mM/s/1%, but that ATP levels would exhaust in approximately a minute if they were limited to 0.079 mM/s/1% (see asterisks in Fig.6E and 6L). When the phosphagen system was removed from the model the volatility of [ATP], its hydrolysis free energy and its production rate increased greatly, but average ATP levels were largely unaffected.

**Figure 6.**
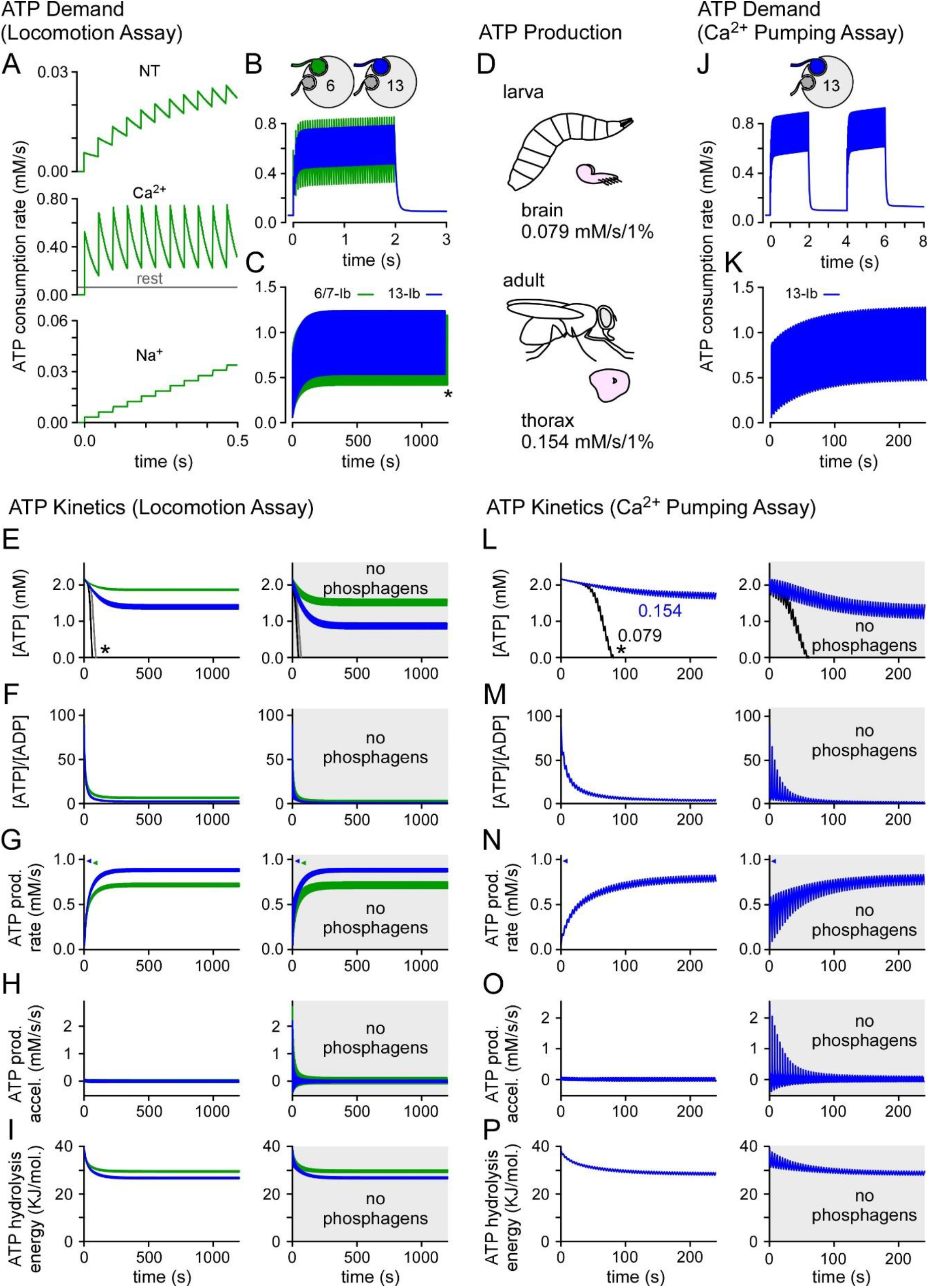
Simulation of ATP levels during prolonged nerve stimulation after ArgK1 KD. **A**. Plots of the ATP consumption rate in response to individual action potentials (APs) in MN6/7-Ib terminals during a peristaltic cycle of 1 s. Three signaling components were considered; SV recycling/refilling associated with neurotransmitter release (NT), Ca^2+^ extrusion and Na^+^ extrusion. A constant value was calculated for all non-signaling activities (the terminal at rest). Units (mM/s) are arrived at by dividing terminal specific ATP consumption rate estimates (ATP molecules/s) by the volume of the terminal. **B**. Plots of the total ATP consumption rate resulting from the summation of each aspect of signaling, along with the non-signaling rate, for two terminals (MN6/7-Ib and MN13-Ib) - representing all type-Ib terminals driven by opsins during the locomotion assay. **C**. Total ATP consumption rate over the 20-minute locomotion assay. The asterisk marks the omission of the final 20 seconds of the MN13-Ib trace to allow terminal comparison. **D**. Estimates of ATP production rates derived from respirometry measurements of isolated larval brains (Neville et al., 2018), and adult thoraces (Jorgensen et al., 2021; Menail et al., 2022). Brains were not otherwise stimulated, but thoraces were permeabilized and stimulated to their maximum rate of O_2_ consumption. Units (mM/s/1%) were arrived at by dividing brain or thorax ATP production rates (mM/s) by tissue specific mitochondrial volume density – yielding mM ATP per second per 1% mitochondrial density. **E–I**. Plots of the cytosolic ATP concentration (E), [ATP]/[ADP] (F), ATP production rate with adult thorax rate caps (0.154 mM/s/1%) denoted on the ordinate (G), ATP production rate acceleration (H) and the free energy available from ATP hydrolysis (I) corresponding to the activity in C, in the presence of a phosphagen system (left panels) and in its absence (right panels). A maximum rate of 0.154 mM/s/1% is used throughout. However, asterisks in E and L indicate where maximum ATP production rate was reduced from 0.154 mM/s/1% to 0.079 [E: MN13-Ib (black line) MN6/7-Ib (grey line); L: MN13-Ib (black line)]. **J**. Plots of the total ATP consumption rate for MN13-Ib terminals - cyclically driven at 50Hz for 2 seconds at 4 second intervals during the Ca^2+^ pumping assay. **K**. Plots of the total ATP consumption rate over the 4 minute period of the Ca^2+^ pumping assay protocol. **L-P**. Plots of the cytosolic ATP concentration (L), [ATP]/[ADP] (M), ATP production rate (N), ATP production rate acceleration (O) and the free energy available from ATP hydrolysis (P) corresponding to the activity in K, in the presence of a phosphagen system (left panels) and in its absence (right panels). The following concentrations and rates were used in E-I and L-P; [ATP] = 2.16 mM, [ADP] = 0.024 mM, [Pi] = 3.8, ATP production limit = 0.154 mM/s/1% (reduced to 0.079 mM/s/1% where indicated by asterisks), K_Ph_ = 39.6, [Arg] = 3.3, [ArgP] = 7.5; sources in Methods.

Consistently, we detected no deficit in the empirical data from the protracted locomotion assay (Fig.5) or the Ca^2+^ pumping assay (Fig.4). Our fluorescent reporters show no indication of an increase in volatility that might betray any increase in volatility in ATP levels.

### Simulations of [ATP] at high firing rates show depletion in the absence of a phosphagen system

The only notable phenotypes in this study occurred when driving MN terminals at firing rates well beyond their endogenous rates. During electrophysiological analyses, we drove firing at 42Hz (Fig.3K-O) and 60Hz (Fig.3B-E), representing multiples of the MN endogenous rates on muscle fiber #6:-2.0X and 2.8X of MN6/7-Ib which fires at an average of 21.4Hz during fictive locomotion, and, 5.4X and 7.7X of MNSNb/d-Is which usually fires at 7.8Hz (Justs et al., 2022). We detected mildly greater depression of neurotransmission over a period of 0.5 seconds at both 60Hz (Fig.3C & E) and 42Hz (Fig.3K & O). Simulation of the 60Hz stimulus protocols (Fig.7A-G) revealed little difference in ATP levels when the phosphagen system was removed from the model (Fig.7C).

However, a stark difference was observed in the ADP/ATP drawdown (Fig.7D), and this was reflected in the reduction in the free energy available from ATP hydrolysis, dropping from 38 to 31 KJ/mol (Fig.7G). An 80Hz stimulation protocol was used in the metabolic imaging protocols (Fig.2), and when simulated here (Fig.7H-N) we observed a substantially greater drawdown in ATP levels (Fig.7J), ADP/ATP ratio (Fig.7K) and the free energy available from ATP hydrolysis (Fig.7N) when the phosphagen system was removed from the model.

### ArgK1 KD leads to deficits in SV exocytosis and recycling

While we are unable to use electrophysiology to examine the impact of ArgK1 KD on neurotransmission during 80Hz stimulation, we can use an optical method if glutamate is added to the bath to stop muscle contraction. SynaptopHluorin is a reporter of SV lumenal pH, and thus reports SV exocytosis as its fluorescence increases when exposed to an extracellular pH near neutral after being quenched at an acidic pH inside SVs (Ng et al., 2002). SynaptopHluorin exhibited bright fluorescence associated with clusters of SVs when expressed in MNs (Fig.7O) and it revealed that net SV exocytosis increased at a slower rate during nerve stimulation when ArgK1 is knocked down (Fig.7P & R; P<0.001) and peaked at a lower value (Fig.7S; P<0.001).

## Discussion

Here we characterized the functional impact of a motor neuron knock down of a phosphagen kinase (ArgK1). We observed mild phenotypes, with impairments in neurotransmission during high-frequency MN firing but we saw no signs of impairment during endurance-related challenges (Table 1). This is reminiscent of phenotypes in muscle-type cytosolic phosphagen kinase (MM-CK) KO mice where contraction force is reduced during tetanus but endurance is relatively unaffected (Dahlstedt et al., 2000; Steeghs, Benders, et al., 1997; van Deursen et al., 1993). Unfortunately, we could find no reports of neurotransmission at the NMJ of CK KO mice that might allow for a comparison with our data, or to determine whether CK KO mice have deficits in neurotransmission concurrent with deficits in muscle physiology. Whether such deficits in neurotransmission would contribute to deficits in musculoskeletal function is unknown as the NMJ safety factor is high in both *Drosophila* and mammals. Never-the-less, the impairments observed here would be expected to have disruptive consequences at central synapses that code information in high firing frequencies. Indeed, mouse KO models of brain-specific CK reveal behavioral and spatial learning deficits (Jost et al., 2002; Streijger et al., 2004; Streijger et al., 2005) and ArgK1 KD flies show deficits in short term memory (Bozzato et al., 2020).

**Table 1.**
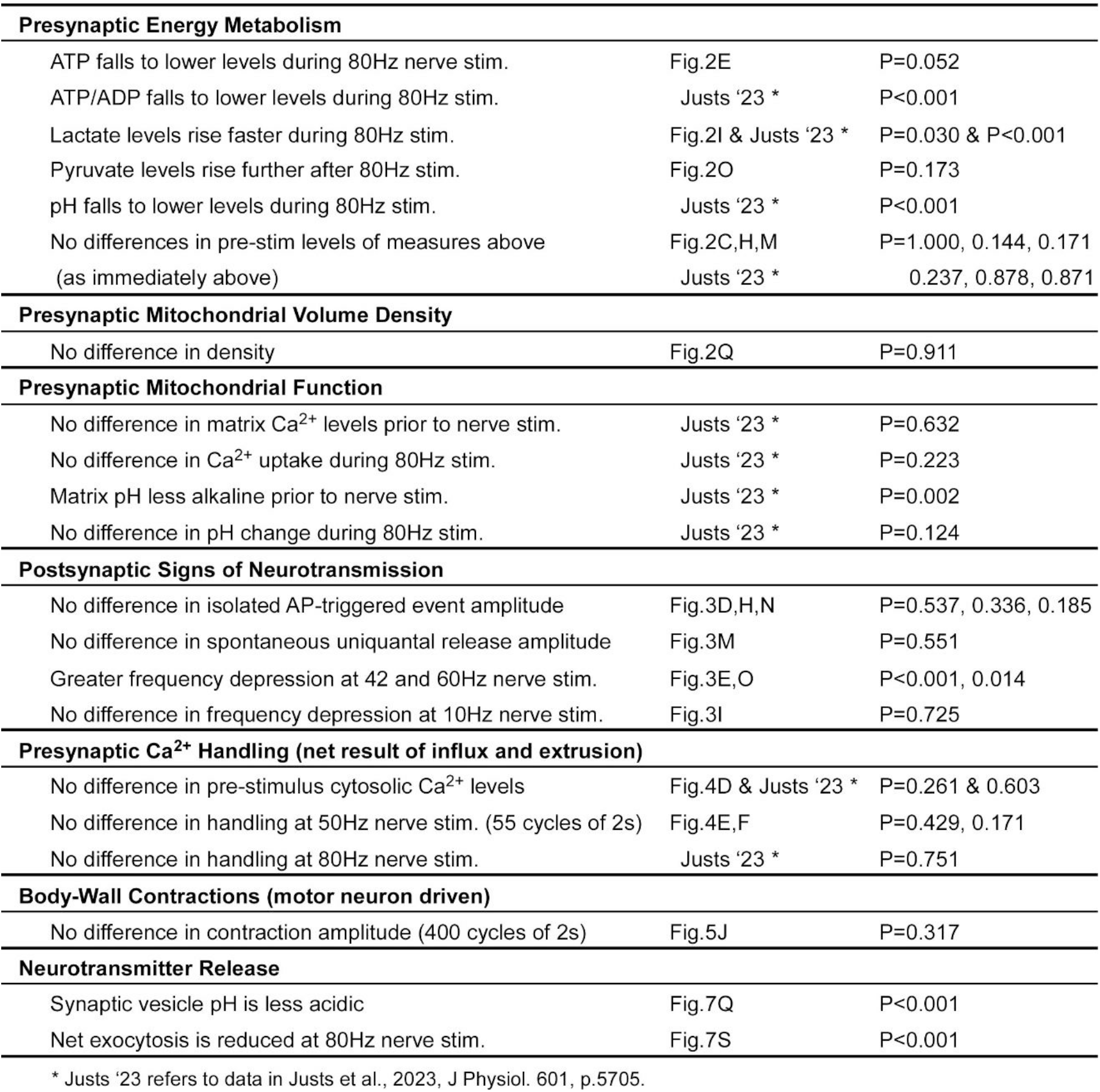
Summary of ArgK1 KD Phenotypes.

The phenotypes observed here mirror findings in MM-CK mice where nerve-evoked contraction force was reduced during tetanus but endurance was relatively unaffected (Steeghs, Benders, et al., 1997). However, as both muscle and nerves were MM-CK deficient, and as the contraction force assays relied on MN function, ambiguity remained regarding the cellular locus of the deficit. This ambiguity was addressed in subsequent experiments, where intact single fibers receiving direct electrical stimulation revealed a deficit in tetanus force during 70Hz stimulation (Dahlstedt et al., 2000). These findings are consistent with deficits we observed in neurotransmission (60Hz, Fig.3E; 42Hz, Fig.3O) and exocytosis (80Hz, Fig.7S). The single fiber studies also used protocols to test endurance but revealed little in the way of deficits stemming from CK KO - similar to our observations of Ca^2+^ handling in MNs (Fig. 4) and sustained body-wall contractions (Fig. 5) in ArgK1 KD *Drosophila* larvae. While it would be desirable to examine mouse MNs and NMJs for deficits concurrent with muscle deficits, the literature leaves no doubt that deficits do exist on the postsynaptic side of the mouse NMJ. Considering the extreme functional differentiation between MNs and muscle fibers, the parallels in the phenotypes are striking.

Tests for concurrent deficits in MN function in CK KO mice were reasonably omitted due to the high “safety factor” at the NMJ, meaning that neurotransmitter release exceeds the threshold required for muscle contraction (Wood & Slater, 2001). With a safety factor of approximately 3-5 fold in mammals and 5-9 fold in *Drosophila* (Marrus & DiAntonio, 2005) it seems unlikely that anything but a profound deficit in MN performance would result in a deficit in contractile force. Ultimately, a muscle contraction force assay (Ormerod et al., 2022) might resolve the capacity of ArgK1 KD MN deficits to impact larval musculoskeletal performance.

Beyond relay synapses like NMJs, the impact of phosphagen systems in neural circuits involving high firing rates is worth considering. Many neurons fire at high rates in the vertebrate central nervous system, with pyramidal neurons, fast-spiking interneurons, and cerebellar mossy fibers capable of firing at 500Hz and above (Delvendahl & Hallermann, 2016). While the cellular locus of the deficit for behavioral and spatial learning deficits has not been elucidated in CK KO mice (Jost et al., 2002; Streijger et al., 2004; Streijger et al., 2005) it was concluded that a lack of CK “rendered the synaptic circuitry in adult brain less efficient in coping with sensory or cognitive activity related challenges” (Streijger et al., 2005). ArgK1 likely fulfils a similar role in central processing in *Drosophila* as ArgK1 KD in the mushroom bodies of adults leads to deficits in short-term memory in a courtship conditioning paradigm (Bozzato et al., 2020).

The ArgK1 KD phenotypes observed at high stimulus frequencies correspond to deficits in ATP levels (Figs 2A-E, 7C and 7J), as well as deficits in the ATP/ADP ratio [Fig.7D and 7K; Justs et al. (2023)] and deficits in the free energy available from ATP hydrolysis (Fig.7G and 7N). The concurrence of these deficits offers little leverage in elucidating whether the phosphagen system primarily acts as a temporal or spatial buffer of adenine nucleotides.

However, as we can detect a deficit in the 60Hz frequency train within 4 impulses (3 intervals at 60Hz = 50ms; Fig. 3C), which is prior to a noticeable deficit in the simulated measures (Fig.7), we propose that the deficits are most pronounced in microdomains that our simulations cannot capture. That is, it is the phosphagen system’s failure to spatial buffer adenine nucleotides that contributes to the ArgK1 KD phenotype. Our simulations represent volume-averaged values, i.e. in the absence of detail regarding the subcellular distribution of ArgK1 we could not build a spatial component. In the cytosolic environment, ADP diffusion is greatly impeded, more so than ATP (Jacobus, 1985; Yoshizaki et al., 1990), and, under conditions of intense ATP hydrolysis, ADP builds up faster than can be cleared by diffusion. In its spatial buffering role, ArgK1 will clear ADP at sites of ATP hydrolysis, offsetting changes in the ATP/ADP ratio and maintaining the free energy of ATP hydrolysis (Mainwood & Rakusan, 1982).

The phenotypes revealed by ArgK1 KD, although mild, can yield some insight into the mechanisms that rely on a presynaptic phosphagen system (Table 1). The greater rate of depression revealed by TEVC (Fig. 3E&O) is an unambiguous indication of a reduced capacity to sustain exocytosis at high firing rates. This conclusion is supported by SynaptopHluorin ArgK1 KD data, where the signal is reduced by half (Fig. 7S). Although changes in SynaptopHluorin fluorescence report exocytosis net of endocytosis, unless we were to propose that endocytosis is accelerated by ArgK1 KD, here too we conclude there is a deficit in exocytosis at high firing rates. This conclusion does not occlude the possibility that the deficit in exocytosis is due to, or in addition to, a deficit in endocytosis. Certainly, the first manifestation of the deficit (≤50ms) would suggest that any deficit in endocytosis imposes its influence in a surprisingly short time frame, synonymous with ultrafast membrane retrieval (Chanaday & Kavalali, 2018; Watanabe et al., 2013).

**Figure 7.**
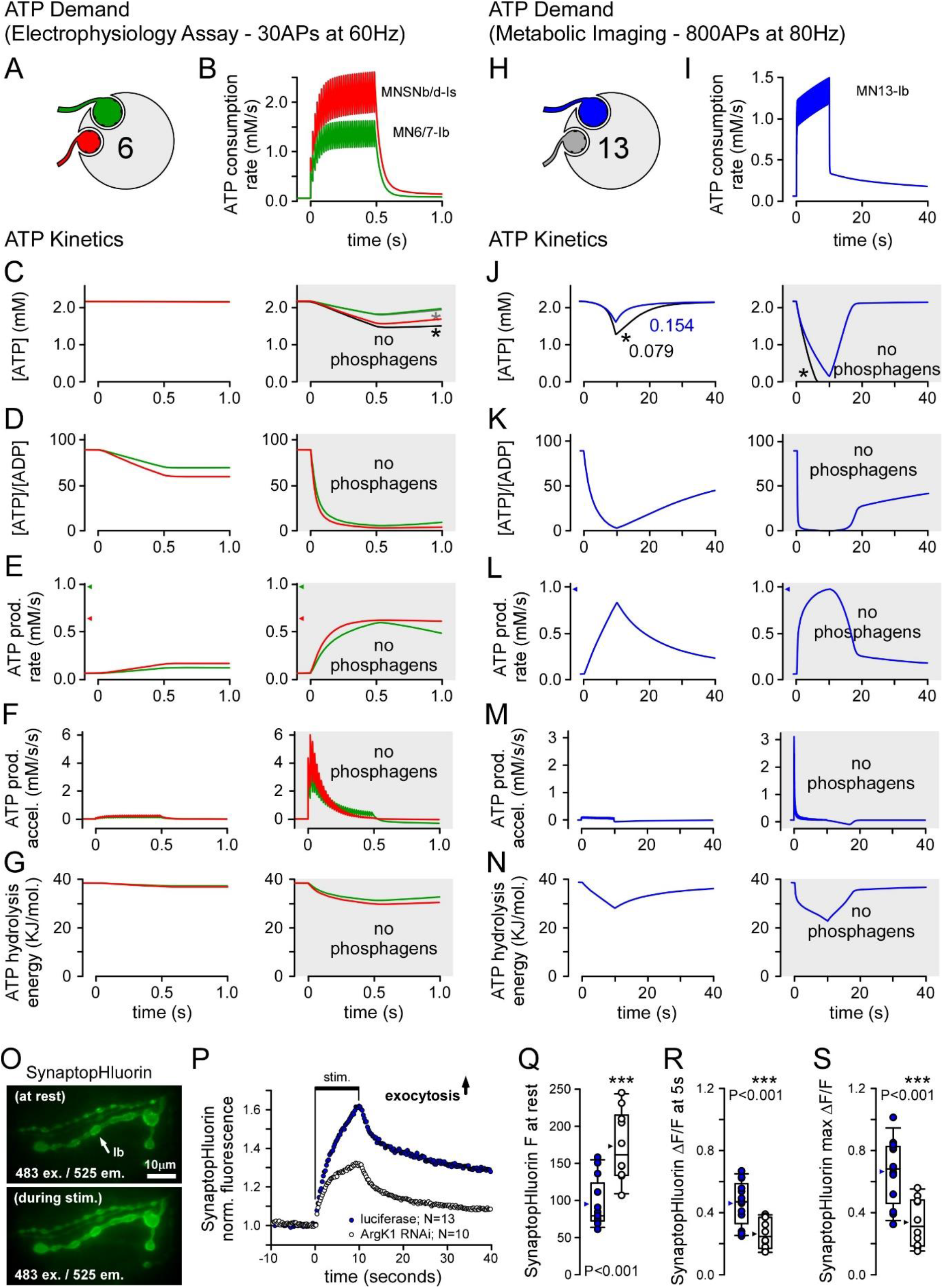
Simulation of ATP levels during high frequency stimulation after ArgK1 KD. **A**. Diagram of the two MN (MN) terminals that release neurotransmitter onto muscle fiber #6 during electrophysiological assays. **B**. Plots of MN terminal ATP consumption rates during a 60Hz train of 30 action potentials (APs). **C–G**. Plots of the cytosolic ATP concentration (C), [ATP]/[ADP] (D), ATP production rate with adult thorax rate caps (0.154 mM/s/1%) denoted on the ordinate (E), ATP production rate acceleration (F) and the free energy available from ATP hydrolysis (G) corresponding to the activity in B, in the presence of a phosphagen system (panels on left) and in its absence (panels on right). A maximum rate of 0.154 mM/s/1% is used throughout. However, asterisks in C and J indicate where maximum ATP production rate was reduced from 0.154 mM/s/1% to 0.079 [C: - Ib (black line) - Is (grey line); J: - Ib (black line)]. **H**. Diagram of the type-Ib MN terminal (MN13-Ib; blue) on muscle fiber #13 that is imaged during metabolic assays. **I**. Plot of MN terminal ATP consumption rate during an 80Hz train of 800 action potentials (APs). **J–N**. Plots of the cytosolic ATP concentration (J), [ATP]/[ADP] (K), ATP production rate (L) ATP production rate acceleration (M) and the free energy available from ATP hydrolysis (N) corresponding to the activity in I, in the presence of a phosphagen system (panels on left) and in its absence (panels on right). The concentrations and rates used in C-G and J-N are the same as those listed in figure 6 with sources given in the Methods. **O**. Images of MN terminals on muscle fiber #13 showing OK6-GAL4 driven expression of exocytosis reporter SynaptopHluorin; captured prior to stimulation (top panel), and during nerve stimulation (bottom panel). **P**. Plots of the normalized average SynaptopHluorin fluorescence in MN13-Ib during nerve stimulation at 80Hz. An increase in the ratio indicates an increase in exocytosis. A measurement was made every 250 ms. **Q**. Plots of Synaptophluorin fluorescence at rest. Each point represents a different preparation. **R**. Change in Synaptophluorin fluorescence (ΔF/F_rest_) after 5 seconds of nerve stimulation. **S**. Greatest change in Synaptophluorin fluorescence (ΔF/F_rest_) after 10 seconds of nerve stimulation. Images and data in O-S were collected from terminals of MN13-Ib in segment 4 in HL6 (2mM CaCl_2_, 15mM MgCl_2_, 7mM L-glutamate). Average traces shown. Box plots show values from all preparations; mean (arrowhead), median (line), 25-75 percentiles box and 10-90% whiskers. Unpaired Student’s t-tests.

Phosphagen kinases colocalize with various ATPases (Schlattner et al., 2016) highlighting mechanisms that might be affected by ArgK1 KD. CK associates both the endoplasmic reticulum and the plasma membrane (Lim et al., 1983; Rossi et al., 1990). Ca^2+^ pumps on both membranes rely on ATP, and there is clear evidence of functional coupling between CK and Ca^2+^ pumps on the endoplasmic reticulum (Korge et al., 1993; Minajeva et al., 1996). However, our previous Ca^2+^ imaging data during high firing rates [800@80Hz; Justs et al. (2023)] show no deficits in Ca^2+^ handling that could explain the exocytosis impairment. Likewise, current data (Fig. 4E) provide no indication of Ca^2+^ handling deficits. CK has also been isolated with cholinergic and glutamatergic synaptic vesicles (SVs) (Friedhoff & Lerner, 1977; Takamori et al., 2006; Wallimann et al., 1985), where it plays a role in SV filling (Xu et al., 1996), but our mEJP amplitude data (Fig. 3K) show no evidence of a filling deficit. Drawing a parallel to CK’s association with myosin in myofibrillar bundles (Krause & Jacobus, 1992), ArgK1 may functionally couple with myosin to transport SVs along presynaptic actin filaments. Actin-myosin-based SV mobilization has been demonstrated at glutamatergic and cholinergic synapses (Mochida, 2020), but whether this process is supported by phosphagen kinases remains unclear. Verstreken et al. (2005) reported that actin-myosin-based mobilization of reserve SVs in *Drosophila* larval MN terminals is vulnerable to low ATP levels, although no acute deficits in exocytosis or endocytosis were detected. Similarly, studies on isolated goldfish retinal bipolar cells (Heidelberger, 2001) and cultured mouse hippocampal neurons (Pathak et al., 2015) found that endocytosis was the first process to fail when ATP levels fell - failing during or before scission. Despite the rapid onset of the deficit (≤50ms) observed in the current study we cannot exclude actin-myosin-based SV mobilization as the primary mechanism affected by ArgK1 KD as knockdown of myosin VI in rat superior cervical ganglion synapses resulted in a deficit within 50ms (Hayashida et al., 2015).

It is possible that the deficit in exocytosis we have observed is due to acidification of the presynaptic compartment rather than a drop in ATP levels. Phosphagen hydrolysis consumes protons and muscle cytosol acidifies more in the absence of a phosphagen system (in ‘t Zandt et al., 1999; Meyer et al., 1986). We previously reported increased activity-dependent acidification of the MN terminal cytosol after ArgK1 KD [P<0.001; Justs et al. (2023)], motivating further analysis of presynaptic metabolism with less pH-sensitive reporters. Several intracellular presynaptic mechanisms are thought to be sensitive to cytosolic acidification, and foremost among these are voltage-gated Ca^2+^ channels (Chesler, 2003). It is known that Ca^2+^ current is inhibited by cytosolic acidification in vertebrate neurons (Mironov & Lux, 1991; Tombaugh & Somjen, 1997), although we observed no deficit in Ca^2+^ handling under the same conditions that evoked greater ArgK1 KD associated acidification [P=0.751; Justs et al. (2023)]. Recent work by Zefirov et al. (2020) showed that intracellular acidification of MN terminals reduced exocytosis possibly due to inhibition of actin-myosin mobilization of synaptic vesicles to release sites.

Acidification is known to effect the kinetics of actin-myosin interactions (Jarvis et al., 2018; Kohler et al., 2012) and the reduction in tetanus force in CK deficient muscle may partially result from the greater rate of acidification in CK deficient muscle (Meyer et al., 1986).

Clarification of the role for phosphagens in MNs could open therapeutic avenues for neurodegenerative diseases involving MN loss and age-related loss of muscle mass and weakness (sarcopenia). Creatine supplementation for these conditions has shown mixed results (Andres et al., 2008; Beal, 2011; Candow et al., 2019; Kreider & Stout, 2021), but there are reasons to be optimistic. First, both neurodegenerative diseases and sarcopenia are heterogeneous groups, and we should expect some sub-groups to be responsive to treatment, while others would be refractory. These sub-groups may reveal themselves as the utility of personalized genetic testing evolves. Secondly, strategies for phosphagen system repair, maintenance or enhancement should not be limited to creatine supplements, but extended to stimulation of the activity and/or expression of CKs and transporters, and the biogenesis of creatine itself. The endogenous regulation of each of these alternative targets is specific to cell type, and a successful strategy for targeting muscle cells might diverge significantly for MNs.

For example, creatine biosynthesis relies on several biosynthetic enzymes and a membrane transporter, and some but not all elements are found in any given cell type, resulting in brain and muscle tissues with different levels of autonomy with regard to creatine biosynthesis (Ensenauer et al., 2004; Joncquel-Chevalier Curt et al., 2015). Similarly, mutations in the creatine transporter gene (SLC6A8) cause different pathologies in brain and muscle (Schoch et al., 2006), highlighting that MNs might respond differently to interventions targeting creatine biogenesis or transporter activity.

## Methods

### 1. Fly stocks

*Drosophila* stocks were raised on standard cornmeal food [Bloomington Drosophila Stock Center (BDSC) recipe] at 22 ± 1°C. Experiments were performed on female 3^rd^ instar larvae with transgenes expressed on a w^1118^ isogenized strain background. BDSC (Bloomington, IN) provided the following fly lines: OK371-GAL4 (#26160), OK6-GAL4 (#64199), nSyb-GAL4 (#51635), UAS-mito-GFP (#8442), ArgK1-GFP (#51522), UAS-ArgK1 dsDNA (#41697), P{CaryP}attP40 (#36304), P{CaryP}attP2 (#36303), UAS-luciferase (#35788), UAS-Dak1 dsDNA (#65107), UAS-Adk2 dsDNA (#55320), UAS-Adk3 dsDNA (#57407) UAS-ChR2(H134R) (#28995). UAS-TagBFP and UAS-SynaptopHluorin were gifts from Dr Kenneth Irvine and Dr Gero Meisenbock, respectively. Lines enabling expression of ATeam1.03^YEMK^, LiLac, Pyronic and oxBFP-tagged ArgK1 exons were made as described below, along with Crispr/Cas9 modified ArgK1 alleles allowing pHusionRed-tagging of the ArgK1 catalytic subunit.

### 2. Genetics

#### Transgenics

GenScript biosynthesized ATeam1.03^YEMK^ DNA (Imamura et al., 2009) which was then subcloned into the MCS of pJFRC14 creating the UAS-ATeam1.03^YEMK^ plasmid. GenScript biosynthesized DNA of LiLac (Koveal et al., 2020) 5’ to DNA of a viral P2A peptide which was 5’ to DNA of Tag-RFP-T, which was then subcloned into the MCS of pJFRC14 creating the UAS-LiLac-P2A-Tag-RFP-T plasmid. Pyronic DNA (San Martin et al., 2014) was acquired from Addgene (#51308) and subcloned into the MCS of pJFPC14 creating the UAS-Pyronic plasmid.

UAS-ATeam1.03^YEMK^ and UAS-LiLac plasmids were injected into *w*^*1118*^ embryos containing the attP40 landing site by Rainbow Transgenic Flies (Newbury Park, CA, USA), and UAS-Pyronic was injected into *w*^*1118*^ embryos containing the attP2 landing site. Strains with transformed germ line cells were balanced and other chromosomes out-crossed to an in-house *w*^1118^ line.

UAS-ArgK-N(exon1)-oxBFP and UAS-ArgK-N(exon1&2)-oxBFP cDNA were synthesized by GenScript and codon optimized for *D. melanogaster*. Both synthesized genes contained BglII and XhoI restriction sites and were cloned into pJFRC-14 plasmids. The UAS-ArgK-N(exon1)-oxBFP construct was designed to have the first exon from ArgK1-RE and ArgK1-RA isoforms (sequence obtained from flybase; 3L: 9064464-9064517) on the N-terminus of oxBFP (Costantini et al., 2015). The UAS-ArgK-N(exon1&2)-oxBFP construct was designed to have the first two exons of the ArgK1-RA isoform (sequence obtained from Flybase; 3L: 9061907-9062467) flanking oxBFP, with the first exon on the N-terminus, and the second exon at the C-terminus (sequence in Supplemental materials, Fig S1). Both constructs were injected into *w*^*1118*^ embryos containing the attP2 landing site by Rainbow Transgenic Flies, then transformants were isolated, balanced and outcrossed as above.

### Targeted genome modification

ArgK1 endogenously tagged FusionRed lines were generated by Genetivision utilizing the two step CRISPR/Cas9-Catalyzed Homology-Directed Repair (HDR) method described by Gratz et al. (2019). The first step replaced part of the endogenous gene with a 3XP3_GFP cassette (which allows easy selection of transformants with eye-specific GFP). The second step replaced the 3XP3-GFP cassette with a donor plasmid carrying one of two mutated ArgK1 alleles.

Step one: nos-Cas9.P embryos (BDSC #54591) were co-injected with a pCFD4 vector (contained gRNA sequences ttctgggtgtagcgagctgaagg & ggctggaggtgcatcgcctccgg) and a donor plasmid. The donor plasmid contained the 3XP3_GFP+ cassette flanked by two 1Kb homologous arms adjacent to a gRNA cleavage site (maps showing the fly genome cleavage sites can be found in the Supplemental materials, Fig. S2). The gRNAs were designed to excise the last exon of all ArgK1 isoforms (which is also the catalytic domain) and the cell’s own endogenous HDR mechanism inserted the 3XP3_GFP+ cassette into the fly’s genome. Flies with the GFP eye-marker were balanced with TM3. This fly line is an ArgK1 knock-out line as the catalytic domain is deleted, and it is homozygous lethal.

Step two: embryos of the nos-Cas9.P;ArgK1 KO(3XP3_GFP+ cassette)/TM3 line were co-injected with pCFD4 vector (containing gRNA sequences gttgtgggatcaagggtgatggg & aatgccgaaggttgccggactgg) and donor plasmid. In this second step, the donor plasmid contained a mutated ArgK1 allele [last exon of ArgK1 endogenously tagged with FusionRed at either the N-terminal or C-terminal (sequence in Supplemental materials, Fig. S2)] flanked by two 1Kb homologous arms adjacent to gRNA cleavage site. The cells own HDR mechanism inserted the mutated allele into the fly’s genome. Individual flies were selected based on loss of GFP eye marker and were balanced with TM3. These lines were then sequenced to determine if the HDR mechanism had integrated ArgK1-FusionRed allele into fly genome. The sequencing results for the FusionRed-N-ArgK1 and FusionRed-C-ArgK1 lines can be found in the Supplemental materials - confirming that ArgK1 was endogenously tagged by FusionRed on the last exon.

#### Knockdown of ArgK1

The effectiveness of ArgK1 KD through presynaptic expression of a dsRNA (BDSC; #41697) directed against ArgK1 message was established previously (Justs et al., 2023), showing near complete knockdown of the ArgK1-GFP signal (98% reduction: P<0.001 by Student’s t test).

### 3. Metabolite, Calcium and SynaptopHluorin Imaging

Five different genetically-encoded functional probes were used in the presynaptic compartment. ATeam1.03^YEMK^ was used to monitor cytosolic ATP levels ([ATP]_c_), LiLac to monitor lactate levels ([lactate]_c_), Pyronic to monitor pyruvate levels ([pyruvate]_c_), mScar8f to monitor Ca^2+^ levels ([Ca^2+^]_c_), and SynaptopHluorin to monitor exocytosis.

Experiments were conducted on female 3^rd^ instar larvae. Filet dissections were performed on a Sylgard tablet in chilled hemolymph-like solution #6 (Macleod et al., 2002) containing CaCl_2_ added to 2mM, and L-glutamic acid added to 7mM [to prevent muscle contraction at 80Hz stimulation; (Macleod et al., 2004)]. Additionally, the cerebral lobes were cut from the ventral ganglia and all nerves connecting the brain to the body muscles were cut, except those to body segment 4. The nerves attached to the ventral ganglion leading to segment 4 were drawn into the lumen of a glass pipette as a loop for subsequent electrical stimulation. Type-Ib motor nerve terminals on muscle 13 of body segment 4 were imaged. High-speed fluorescence microscopy was performed on a Nikon Eclipse FN1 microscope fitted with a Nikon 100X water-immersion (1.1 NA) objective. Probes were excited sequentially by a Lumencor SPECTRA-X light source. Emitted light was captured by two Andor iXon3 EMCCD cameras (DU-897) mounted on an Andor TuCam beam splitter (Andor Technology, South Windsor, CT). Filters and dichroic mirrors were obtained from Chroma Technology (Bellows

Falls, VT, USA) or Semrock (Lake Forest, IL, USA). ATeam1.03^YEMK^, LiLac with TagRFP-T, and Pyronic were all excited by reflection off a triple-edge dichroic mirror (Chroma, 69008bs), with emitted light passing back through the dichroic and a 470/24, 535/30 and 632/60nm triple-band emission filter (Chroma, 69008bs). mScar8f and SynaptopHluorin was excited by reflection off a quadruple-edge dichroic mirror (Chroma, 89100bs), with emitted light passing back through the dichroic and a 455/50, 525/36, 605/52 and 705nm/72nm quad-band emission filter. Nikon Elements software controlled the illumination sequence and camera image acquisition. A Master 8 (A.M.P.I., Israel) controlled the timing of nerve stimulation via an A-M Systems Model 2100 Isolated Pulse Stimulator. Terminals were allowed to equilibrate for at least 2 minutes after any nerve stimulus train.

To monitor changes in [ATP]_c_ we compared the fluorescence emitted from CFP with mVenus of the ATeam1.03^YEMK^ construct. CFP of ATeam1.03^YEMK^ was excited at 434/17nm, and emitted light was reflected by a second dichroic mirror (509nm edge) before being collected to a camera through a 475/28nm filter. ATeam1.03^YEMK^ was also excited by 434/17nm light, and mVenus emission was collected by a second camera after passing through the second dichroic and a 525/30nm filter.

To monitor changes in [lactate]_c_ we compared the fluorescence emitted from mTurquoise2 of LiLac with TagRFP-T. LiLac was excited at 436/20nm and a camera collected the emitted light, while TagRFP-T was excited by 510/25nm light, and its emission was collected by the same camera.

To monitor changes in [pyruvate]_c_ we compared the fluorescence emitted from mTFP with Venus, both within the Pyronic construct. mTFP was excited at 434/17nm, and emitted light was reflected by a second dichroic mirror (509nm edge) before being collected to one camera through a 475/28nm filter. Pyronic was also excited by 434/17nm light, and Venus emission was collected by a second camera after passing through the second dichroic and a 525/30nm filter.

To monitor changes in [Ca^2+^]_c_ we compared the fluorescence emitted from Ca^2+^-sensitive GCaMP8f with Ca^2+^-insensitive mScarlet, both within the mScar8f construct. GCaMP8f was excited at 483/32nm, and emitted light was reflected by a second dichroic mirror (580nm edge) before being collected to a camera through a 525/50nm filter. mScarlet was excited by 550/25nm light, and its emission was collected by a second camera after passing through the second dichroic and a 617/73nm filter.

For each of ATeam1.03^YEMK^, LiLac and Pyronic, 4 images pairs were collected every second - yielding 4 fluorescence ratio estimates. Ten (10) image pairs were collected for mScar8f. The ratio of intensities, relative to the ratio prior to nerve stimulation, was plotted against time as ΔR/R. The ATeam1.03^YEMK^ fluorescence ratio was corrected for bleaching by fitting a monoexponential to 20 seconds of pre-stimulus ratio estimates and using this fit to correct the ratio between time=0 and time=40 in Fig.2B, e.g. Macleod (2012).

SynaptopHluorin was used to monitor changes in exocytosis. It uses SV membrane protein Synaptobrevin to target SuperEcliptic-pHluorin (Sankaranarayanan et al., 2000) to the lumen of SVs (Ng et al., 2002). SynaptopHluorin was the only functional reporter not used in a ratiometric mode. It was excited with 483/32nm light reflected by a quad-band dichroic mirror, and emitted light passed back through the dichroic and a 525/50nm emission filter and collected by a single camera. Fluorescence intensity estimates (F), relative to the fluorescence prior to nerve stimulation, was plotted against time as ΔF/F.

Fluorescence intensity measurements were performed on ImageJ. Movement during imaging was corrected using the Template Matching plugin by aligning consecutive frames. ROIs covered non-distal boutons along a length of 20 µm of the nerve terminal, with two background ROIs for reference. The average pixel intensity values were calculated in each channel for each ROI for each frame and exported to Microsoft Excel. The average of the two background ROIs was subtracted from the average bouton ROI in each channel for each frame. Preparations were not used if the boutons exhibited movement out of frame or out of focus.

Outliers were assessed using the median absolute deviation [MAD; Leys et al. (2013)] of the baseline and delta measurements, where an outlier was any value beyond either the median +3 x MAD or −3 x MAD.

### 4. Live Imaging of ArgK1-GFP, -oxBFP and -FusionRed

All imaging was conducted on female 3^rd^ instar larvae filet-dissected on a Sylgard tablet in chilled HL6 with 2mM CaCl_2_. Preparations were then rinsed thrice in 0.1 mM CaCl_2_ HL6, covered with a glass coverslip, and imaged with a Nikon 60X, 1.20 NA, Plan Apochromat VC water-immersion objective on a Nikon A1R CLSM fitted with GaAsP detectors. Sequentially, scanning was used, starting with the longest laser wavelengths and progressing to the shortest (560nm, 488nm, 405nm). Images represent a collapsed Z-series encompassing a limited depth of the ventral ganglion or the full depth of terminal boutons in the periphery (3 μm and 1 μm step sizes, respectively).

### 5. Electrophysiology

Electrophysiology was exclusively conducted on female 3^rd^ instar larvae in either hemolymph-like solution #3 [HL3; Stewart et al. (1994)] or HL6 (Macleod et al., 2002) containing added MgCl_2_ and CaCl_2_ as described in figure 3. Fillet dissections were performed in chilled HL6 on Sylgard tablets and recordings were made 20 to 60 minutes after transecting the segmental nerves. Signals were detected, digitized and recorded using an Axoclamp 900A amplifier (Molecular Devices; Sunnyvale, CA) connected to a 4/35 PowerLab (ADInstruments; Colorado Springs, CO) and a PC running LabChart v8.0. Micropipettes filled with a 1:1 mixture of 3 M KCl and 3 M K-acetate. Measurements were performed on segment #4 under a 20X water-dipping objective of a BX50WI Olympus microscope to allow unequivocal identification of muscle fibers. Two-electrode voltage clamp (TEVC; clamped to −70mV) was used to quantify the combined release from MN6/7-Ib and MNSNb/d-Is onto body-wall muscle fiber #6. A suction pipette applied 0.3ms electrical impulses to the transected nerve to evoke release from both MN terminals. The voltage used was 20-50% above the threshold needed to initiate an AP in both MNs. Preparations were discarded if we observed a failure to evoke release from both MNs in response to any AP in a stimulus train, or if the unclamped resting membrane potential (RMP) dropped below −55 mV. Where Quantal Content (QC) was calculated (Fig 3H & 3N), a minimum of 10 Excitatory Junctional Potentials (EJPs) were recorded during 0.2Hz stimulation, along with 30 miniature EJPs (mEJPs), prior to implementing a TEVC. QC was calculated by dividing the corrected mean EJP amplitude by the mean mEJP amplitude. Mean EJP amplitude was corrected for non-linear summation (McLachlan, 1981). The median absolute deviation (MAD) was used to assess outliers as described in figure 3.

### 6. In Vivo MN Performance Assay

We have described assembly of the apparatus to evoke body-wall contractions in *Drosophila* larvae, preparation of larvae for this assay, and data analysis in a separate Methods article (Arab et al., 2025). Briefly, we expressed a light-activated opsin (H134R-ChR2) in motor-neurons using OK371-GAL4, raised the larvae in the dark on food supplemented with all-trans retinal (100mg/ml) and drove musculoskeletal contractions using blue light in an illumination cycle of 2 seconds on, and 1 second off. Images of the larvae collected on a CCD camera at a rate of 10 frames-per-second were analyzed offline using custom-built software.

### 7. Statistical analysis

Student’s t-tests were performed to determine significance in individual comparisons. The Mann-Whitney Rank Sum test was applied where samples failed a normality test. One way ANOVA was used for multiple comparisons followed by post-hoc tests.

### 8. Simulation of ATP Dynamics

The simulation model is described in Justs et al. (2023). The only differences are the timing of action potentials, described in the main text, and the initial values of [P_i_] and [ADP], discussed in “Parameter Values” below. The model tracks the concentrations of six metabolites ([ATP], [ADP], [AMP], [ArgP], [Arg] and [P_i_]) that change according to three processes – ATP Consumption, ATP Production and Phosphagen Equilibration.

#### ATP consumption

ATP consumption is represented by the time rate of [ATP] reduction, in exchange for [ADP] + [P_i_].

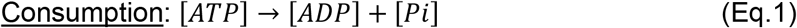

Action potential (*AP*)-associated ATP consumption processes (signaling processes) are modeled as an instantaneous spike, followed by an exponential decay over the time associated with each of the ATP consumption processes; Na^+^ extrusion, Ca^2+^ extrusion and SV recycling/refilling associated with neurotransmitter release (NT). That is, each process j will consume Nj ATP molecules, decay with a time constant τj, and manifest a spike of height Nj / τj at the time of firing. Meanwhile, a static base rate of ATP consumption due to non-signaling activities, unrelated to APs, is applied throughout the time of simulation. The time evolution of ATP consumption from a single AP occurring at time *t*_*0*_ is then:

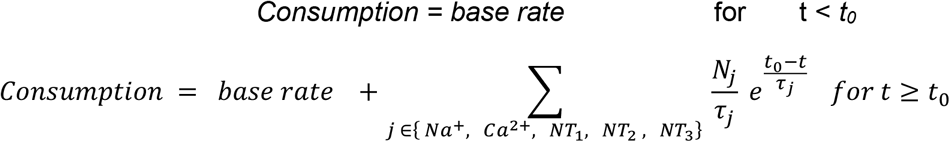

During a train with a firing rate (FR; Hz), a new spike occurs every 1/FR seconds. Each new spike summates upon the transients of previous spikes. A *full cycle* is the span of time from the beginning of one train of spikes to the beginning of the next train, which corresponds to a contraction cycle during locomotion. The *duty cycle* is the amount of time during a *full cycle* when a train is firing. As an example, for a train of APs firing at 10Hz, starting at t=t_0_, with a *full cycle* of 1 second and *duty cycle* of 0.6 seconds, 7 spikes will occur, 100ms apart, from time t_0_ to t_0_ + 0.6 seconds. The remaining time of the full cycle will not incur another spike, and another train with the same parameters will start at t_0_ + 1 seconds.

#### ATP production

ATP production is represented by the time rate of the decrease in [ADP] and [P_i_], in exchange for an increase in [ATP].

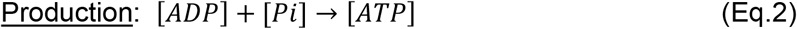

We utilize the ATP production model from (Wilson, 2017). A production cap, representing the maximal rate of ATP production set by the available mitochondrial volume within the terminal, restricts the ATP production rate. An interpolation function passes the production rate curve to the production cap smoothly if ATP production approaches its cap.

#### Equilibration

Adenylate Kinase and Arginine Kinase maintain the following equilibria at all times:

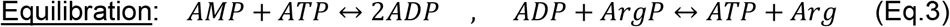

#### Numerical Integration

At each time step, [ADP] and [ATP] are updated first using the production and consumption rates. Then, [AMP], [ADP], [ATP], [Arg] and [ArgP] are updated according to (Eq.3) until their equilibrium relations, below, are satisfied

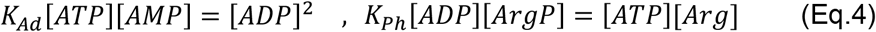

where K_Ad_ and K_Ph_ are the equilibrium constants for Adenylate Kinase and Arginine Kinase, respectively. When simulating without phosphagen, the 2^nd^ equation in (Eq.4) is not enforced.

#### Parameter Values

Following Justs et al. (2023), we use the most complete set of model parameter values from the nervous system of a single invertebrate we know of: the giant squid axon. Initial concentrations are set to 2.16 mM [ATP] (Caldwell, 1960; Mullins & Brinley, 1967; Requena et al., 1979), 7.5 mM [ArgP] (DiPolo & Beauge, 1996), and 3.3 mM [Arg] (Deffner, 1961; Deffner & Hafter, 1959). K_Ph_ is set to 39.6 (Teague & Dobson, 1999). K_Ad_ is set to 1 (Wilson, 2017). [AMP] and [ADP] initial concentrations are derived from (Eq.4) using the above values for [ATP], [ArgP], [Arg], K_Ph_ and K_Ad_. Squid axon [Pi] measurements range from 3.8 to 17.8 mM (Caldwell, 1960; Deffner, 1961). To ensure the fairest comparison between systems with and without a phosphagen system, the resulting values (24.2 mM [ADP] and 0.27 mM [AMP]) are used whether or not a phosphagen system is present. Since ionic concentrations in terrestrial insect nerve tissue are uniformly lower than in their marine invertebrate counterparts [*Periplaneta* and *Romalea*; Potts and Parry (1964)], we now use the lower end of the squid axon range: [Pi]=3.8mM. This choice is further supported by comparison with the terrestrial *Manduca Sexta* P concentrations, where tissue [Pi] ranges from 1.2 - 3.8 mM (Woods et al., 2002).

## Acknowledgements

This work was supported by NIH NINDS awards NS103906 and NS123377 to Dr. Gregory Macleod. Dr. Karlis Justs’ current affiliation is with Agilent Technologies USA. Mr. Carlos Oliva’s current affiliation is with the University of Miami. Mr. Gabriel Bonassi’s current affiliation is with the University of Toledo College of Medicine and Life Sciences. We are grateful for discussions with Drs Ronald Lynch and Håkan Westerblad. Stocks obtained from the Bloomington Drosophila Stock Center (BDSC: NIH P40OD018537) were used in this study.

