## Supplemental materials for "Optimal Neuromuscular Performance Requires Motor Neuron Phosphagen Kinases"

**Supplemental Figure 1.**

Annotated cDNA sequences for ArgK-N(exon1)-oxBFP and ArgK-N(exon1&2)-oxBFP constructs prior to cloning into pJFRC14 plasmids and the generation of transgenic animals.

Color code: BglII restriction site Kozak Sequence Linker N-terminal of ArgK1-RE and ArgK1-RA Isoforms (exon1) Second exon from N-terminal of Argk1-RA Isoform oxBFP Stop Codon XhoI restriction site Extra nucleotide to keep protein in frame

ArgK-N(exon1)-oxBFP:

AGATCTCAAGATGTTCGCCCTGTGGTACCTGACCTTCGCCGTGGATGAGATTCGCGGCGGCGGAAG
CGGCGGCGGCAGCATGGTGTCCAAGGGAGAGGAGCTGTTCACCGGAGTGGTGCCCATCCTGGTGG
AGCTGGATGGCGACGTGAATGGACACAAGTTCTCCGTGCGCGGCGAGGGAGAGGGCGACGCCACC
AACGGCAAGCTGACCCTGAAGTTCATCTCGACCACCGGCAAGCTGCCGGTGCCCTGGCCCACCCTG
GTGACCACCCTGTCGCACGGAGTGCAGGTGTTCGCCCGCTACCCGGATCACATGAAGCAGCACGA
CTTCTTCAAGTCCGCCATGCCGGAGGGATACGTGCAGGAGCGCACCATCTTCTTCAAGGATGACGG
CACCTACAAGACCCGCGCCGAGGTGAAGTTCGAGGGCGATACCCTGGTGAACCGCATCGAGCTGA
AGGGCGTGGATTTCAAGGAAGATGGCAATATCCTGGGCCACAAGCTGGAGTACAACTTCAATAGCC
ACAACATCTACATCATGGCCGTGAAGCAGAAGAACGGCATCAAGGCCAATTTCAAGATCCGCCACAA
TGTGGAGGATGGCTCCGTGCAGCTGGCCGACCACTACCAGCAGAACACCCCCATTGGCGATGGAC
CCGTGCTGCTGCCGGACTCCCACTACCTGTCGACCCAGAGCGTGCTGTCCAAGGATCCCAATGAGA
AGCGCGACCACATGGTGCTGCTGGAGTTCCGCACCGCCGCCGGAATCACCCTGGGCATGGACGAG
CTGTACAAGTAATAACTCGAG

ArgK-N(exon1&2)-oxBFP:

AGATCTCAAGATGTTCGCCCTGTGGTACCTGACCTTCGCCGTGGATGAGATTCGCGGCGGCGGAAG
CGGCGGCGGCTCCATGGTGTCGAAGGGAGAGGAGCTGTTCACCGGAGTGGTGCCAATCCTGGTGG
AGCTGGATGGCGACGTGAATGGACACAAGTTCTCGGTGCGCGGCGAGGGAGAGGGCGACGCCACC
AACGGCAAGCTGACCCTGAAGTTCATCTCCACCACCGGCAAGCTGCCCGTGCCCTGGCCCACCCTG
GTGACCACCCTGTCCCACGGAGTGCAGGTGTTCGCCCGCTACCCCGATCACATGAAGCAGCACGAC
TTCTTCAAGTCGGCCATGCCCGAGGGATACGTGCAGGAGCGCACCATCTTCTTCAAGGATGACGGC
ACCTACAAGACCCGCGCCGAGGTGAAGTTCGAGGGCGATACCCTGGTGAACCGCATCGAGCTGAA
GGGCGTGGATTTCAAGGAAGATGGCAATATCCTGGGCCACAAGCTGGAGTACAACTTCAATTCCCAC
AACATCTACATCATGGCCGTGAAGCAGAAGAACGGCATCAAGGCCAATTTCAAGATCCGCCACAATG
TGGAGGATGGCAGCGTGCAGCTGGCCGACCACTACCAGCAGAACACCCCGATTGGCGATGGACCG
GTGCTGCTGCCCGACTCCCACTACCTGAGCACCCAGTCCGTGCTGTCGAAGGATCCGAATGAGAAG
CGCGACCACATGGTGCTGCTGGAGTTCCGCACCGCCGCCGGAATCACCCTGGGAATGGATGAGCT
GTACAAGGGCGGCGGCAGCGGCGGCGGCAGCCGCAAGCGCCTGGCCTGGCTGTTCAGCTCCAAC
AAGCCGGCCGCCCCCGCCCTGGACAACAAGCCGGCCAATCCGGCCCCCGCCAAGGAGTCCGCCC
CGGCCCCCGCCCCGGCCCCCACCCCCAAGCCGGCCGTGGTGCCCCCGGCCCCGAAGCCGGATCC
CCCGAAGCCCGCCCCGGCCGTGGCCAAGCCGACCCCGGTGCCCGTGCCGGCCACCGCCCCGGCC
CCACCCAAGGAAGAGCCCGCCCCGAAGCCCAAGCCCGAGCCGGTGCCCTCCCCGGTGGTGGCCC
CACCCAAGCCCACCCCTCCGCCAGCCAAGCCGTCGAGCCCGCCCAAGCAGGCCGACAAGATGCCG
ATCCCCCTGCCGAAGAGCCTGACCGAGGCCAACACCAATGGCCAGAACGGCAATGCCGCCAACGG
CGGAAATGTGGATGAGCTGGTGTTCGGCGGACAGCAGGCCGAGAAGGTGCTGCCGGCCGCCAAGG
AAGCCTCGAACGACTTCATCAAGGGCGAGACCAATGCCTTCATCCAGAGCATCAAGGAAGCCCAGC
AGCTGGGCGAGCGCTAATAACTCGAG

**Supplemental Figure 2.**

Genomic Map Sequences for wildtype, FusionRed-N-ArgK1 and FusionRed-C-ArgK1 fly lines (only in region around last major exon):

Color Code: Intron ArgK1 Last Exon FusionRed Linker gRNA pair (first step) gRNA pair (second step)

Genomic Sequence of wildtype *D. melanogaster* on Region 3L: compliment 9,064,441-9,065,883 (obtained from flybase.org)

tcagtcagccaccaccgaaagcaggacacaatcgagaaacaaaaccagagtcaccaccactaaaaaaggggcgttgtgggatcaagggtga
tgggtgttcggttctgggtactgcttctgggtgtagcgagctgaagggaaagggagcgtgggtatatgtggcagtcgcgtgccacagacccaagt
cgagtatgagcacatggccataacccgaaactaattgcggggcaaacaggcaggaattgaggaaaacagaaacacaagaaaacatgttgccg
acttttgggcgccacaaaatgaagataggcgagtcctttgtagcaataaaatgttgtactgtcgccccagccacgcccttttactaatggtcaataccc
cattctctccttctctctacgtatttttgcagCAAACAAGACACCATGGTTGATGCCGCTGTTCTCGCTAAACTGGAGGA
GGGTTATGCCAAGTTGGCTGCCTCCGACTCCAAGTCGCTGTTGAAGAAGTACCTGACCAAGGAGGT
CTTCGACAACCTGAAGAACAAGGTCACGCCCACCTTCAAGTCGACCCTGCTGGATGTGATCCAGTCT
GGCCTGGAGAACCACGATTCCGGCGTCGGCATCTACGCTCCCGATGCCGAGGCTTACACAGTGTTC
GCCGACCTGTTCGATCCCATCATCGAGGACTACCATGGTGGCTTCAAGAAGACCGACAAGCACCCG
GCCTCCAACTTTGGCGATGTGTCCACCTTCGGCAACGTTGACCCCACCAACGAGTACGTGATCTCCA
CTCGCGTGCGTTGCGGTCGCTCCATGCAGGGATACCCCTTCAACCCCTGCTTGACCGAGGCCCAGT
ACAAGGAGATGGAAAGCAAGGTCAGCAGCACCCTGTCCGGTCTGGAAGGTGAGCTGAAGGGCAAG
TTCTACCCCCTGACTGGCATGGAGAAGGCCGTCCAGCAGCAGCTGATCGACGACCACTTCCTGTTC
AAGGAGGGCGATCGTTTCCTGCAGGCCGCCAACGCCTGCCGCTTCTGGCCCAGCGGCCGTGGCAT
CTACCACAACGATGCCAAGACCTTCCTGGTCTGGTGCAACGAGGAGGACCATCTCCGCATCATCTC
CATGCAGCAGGGTGGTGATCTGGGCCAGATCTACAAGCGTCTGGTGACCGCCGTCAACGAAATCGA
GAAGCGTGTGCCATTCAGCCACGACGACCGTCTCGGTTTCCTGACCTTCTGCCCCACCAACCTGGG
CACCACCATCCGTGCCTCCGTGCACATCAAGGTGCCCAAGCTGGCATCCAACAAGGCCAAGCTGGA
GGAGGTTGCCGCCAAGTACAACCTGCAGGTGCGCGGAACCCGCGGTGAGCACACCGAGGCTGAGG
GTGGTGTCTACGACATCTCCAACAAGCGTCGCATGGGTCTGACCGAGTTCGAGGCCGTCAAGGAGA
TGTACGATGGCATCACCGAGCTGATCAAGCTCGAGAAGAGCCTGTAAATTGCCCGCCAACCGCCGA
GCATCGTCCCCCCGGCTGGAGGTGCATCGCCTCCGGAGTTAACATCTATCCACGGGTGCCACCAGT
TCGGGGCATACGCAACAATCAAGAACAACAACAGCAACACTTAGAAATCTTCGGCTCGTCCGCCTTG
TCGCTTTTGGGCGGTTCGAACTGCAAATATGTCTAGTTTGTAGCTATGGTGTGGAGCTGCTGTCCCA
GTCCGGCAACCTTCGGCATTTTCGTATACTTTGAGGGTCGCGGGCGCAGGCAGGGATCCCATCTTG
CAACACCATCAACACCTACAGCAGCAGCAGCAACAACGGATAACTATTTGACAAAAACATTTGCTT

Genomic Sequence of FusionRed-N-ArgK1 line (only last major exon is shown):

ttttgcagCAAACAAGACACCATGGGTGGCAGTGGTGGCTCGGTGAGCGAGCTGATTAAGGAGAACATG
CCCATGAAGCTGTACATGGAGGGCACCGTGAACAACCACCACTTCAAGTGCACATCCGAGGGCGAA
GGCAAGCCCTACGAGGGCACCCAGACCATGAGAATCAAGGTCGTCGAGGGCGGCCCTCTCCCCTT
CGCCTTCGACATCCTGGCTACCAGCTTCATGTACGGCAGCAGAACCTTCATCAAGCACCCTCCGGG
CATCCCCGACTTCTTTAAGCAGTCCTTCCCTGAGGGCTTCACATGGGAGAGAGTCACCACATACGAA
GACGGGGGCGTGCTGACCGCTACCCAGGACACCAGCCTCCAGGACGGCTGCCTCATCTACAACGT
CAAGGTTAGAGGGGTGAACTTCCCAGCCAACGGCCCTGTGATGCAGAAGAAAACACTCGGCTGGGA
GGCCTCCACCGAGACGATGTACCCCGCTGACGGCGGCCTGGAAGGCGCATGTGACATGGCCCTGA
AGCTCGTGGGCGGGGGCCACCTGATCTGCAACCTTGAGACCACATACAGATCCAAGAAACCCGCTA
CGAACCTCAAGATGCCCGGCGTCTACAACGTGGACCACAGACTGGAAAGAATCAAGGAGGCCGACG
ATGAGACCTACGTCGAGCAGCACGAGGTGGCTGTGGCCAGATACTCTACTGGTGGCGCTGGTGATG
GAGGTAAAGGTGGCTCTGGCGGTAGTGTTGATGCCGCTGTTCTCGCTAAACTGGAGGAGGGTTATG
CCAAGCTGGCTGCCTCCGACTCCAAGTCGCTGTTGAAGAAGTACCTGACCAAGGAGGTCTTCGACA

ACCTGAAGAACAAGGTCACGCCCACCTTCAAGTCGACCCTGCTGGATGTGATCCAGTCTGGCCTGG
AGAACCACGATTCCGGCGTGGGCATCTACGCTCCCGATGCCGAGGCTTACACAGTGTTCGCCGACC
TGTTCGACCCCATCATCGAGGACTACCATGGTGGCTTCAAGAAGACCGACAAGCACCCGGCCTCCA
ACTTTGGCGATGTGTCCACCTTCGGCAACGTTGACCCCACCAACGAGTACGTGATCTCCACTCGCGT
GCGTTGCGGTCGCTCCATGCAGGGATACCCCTTCAACCCCTGCTTGACCGAGGCCCAGTACAAGGA
GATGGAAAGCAAGGTCAGCAGCACCCTGTCCGGTCTGGAAGGTGAGCTGAAGGGCAAGTTCTACCC
CCTGACTGGCATGGAGAAGGCCGTCCAGCAGCAGCTGATCGACGACCACTTCCTGTTCAAGGAGGG
CGATCGTTTCCTGCAGGCCGCCAACGCCTGCCGCTTCTGGCCCAGCGGCCGTGGCATCTACCACAA
CGATGCCAAGACCTTCCTGGTCTGGTGCAACGAGGAGGACCATCTCCGCATCATCTCCATGCAGCA
GGGTGGTGATCTGGGCCAGATCTACAAGCGTCTGGTGACCGCCGTCAACGAAATCGAGAAGCGTGT
GCCATTCAGCCACGACGACCGTCTCGGTTTCCTGACCTTCTGCCCCACCAACCTGGGCACCACCAT
CCGTGCCTCCGTGCACATCAAGGTGCCCAAGCTGGCATCCAACAAGGCCAAGCTGGAGGAGGTTG
CCGCCAAGTACAACCTGCAGGTGCGCGGAACCCGCGGTGAGCACACCGAGGCTGAGGGTGGTGTC
TACGACATCTCCAACAAGCGTCGCATGGGTCTGACCGAGTTCGAGGCCGTCAAGGAGATGTACGAT
GGCATCACCGAGCTGATCAAGCTCGAGAAGAGCCTGTAAATTGCCCGCCAACCGCCGAGCATCGTC
CCCCC

Genomic Sequence of FusionRed-C-ArgK1 line (only last major exon is shown):

ttttgcagCAAACAAGACACCATGGTTGATGCCGCTGTTCTCGCTAAACTGGAGGAGGGTTATGCCAAGC
TGGCTGCCTCCGACTCCAAGTCGCTGTTGAAGAAGTACCTGACCAAGGAGGTCTTCGACAACCTGA
AGAACAAGGTCACGCCCACCTTCAAGTCGACCCTGCTGGATGTGATCCAGTCTGGCCTGGAGAACC
ACGATTCCGGCGTGGGCATCTACGCTCCCGATGCCGAGGCTTACACAGTGTTCGCCGACCTGTTCG
ACCCCATCATCGAGGACTACCATGGTGGCTTCAAGAAGACCGACAAGCACCCGGCCTCCAACTTTG
GCGATGTGTCCACCTTCGGCAACGTTGACCCCACCAACGAGTACGTGATCTCCACTCGCGTGCGTT
GCGGTCGCTCCATGCAGGGATACCCCTTCAACCCCTGCTTGACCGAGGCCCAGTACAAGGAGATGG
AAGGCAAGGTCAGCAGCACCCTGTCCGGTCTGGAAGGTGAGCTGAAGGGCAAGTTCTACCCCCTGA
CTGGCATGGAGAAGGCCGTCCAGCAGCAGCTGATCGACGACCACTTCCTGTTCAAGGAGGGCGATC
GTTTCCTGCAGGCCGCCAACGCCTGCCGCTTCTGGCCCAGCGGCCGTGGCATCTACCACAACGATG
CCAAGACCTTCCTGGTCTGGTGCAACGAGGAGGACCATCTCCGCATCATCTCCATGCAGCAGGGTG
GTGATCTGGGCCAGATCTACAAGCGTCTGGTGACCGCCGTCAACGAAATCGAGAAGCGTGTGCCAT
TCAGCCACGACGACCGTCTCGGTTTCCTGACCTTCTGCCCCACCAACCTGGGCACCACCATCCGTG
CCTCCGTGCACATCAAGGTGCCCAAGCTGGCATCCAACAAGGCCAAGCTGGAGGAGGTTGCCGCC
AAGTACAACCTGCAGGTGCGCGGAACCCGCGGTGAGCACACCGAGGCTGAGGGTGGTGTCTACGA
CATCTCCAACAAGCGTCGCATGGGTCTGACCGAGTTCGAGGCCGTCAAGGAGATGTACGATGGCAT
CACCGAGCTGATCAAGCTCGAGAAGAGCCTGGGTGGCTCTGGCGGTAGTGGTGGCAGTGGTGGCT
CGGTGAGCGAGCTGATTAAGGAGAACATGCCCATGAAGCTGTACATGGAGGGCACCGTGAACAACC
ACCACTTCAAGTGCACATCCGAGGGCGAAGGCAAGCCCTACGAGGGCACCCAGACCATGAGAATCA
AGGTCGTCGAGGGCGGCCCTCTCCCCTTCGCCTTCGACATCCTGGCTACCAGCTTCATGTACGGCA
GCAGAACCTTCATCAAGCACCCTCCGGGCATCCCCGACTTCTTTAAGCAGTCCTTCCCTGAGGGCTT
CACATGGGAGAGAGTCACCACATACGAAGACGGGGGCGTGCTGACCGCTACCCAGGACACCAGCC
TCCAGGACGGCTGCCTCATCTACAACGTCAAGGTTAGAGGGGTGAACTTCCCAGCCAACGGCCCTG
TGATGCAGAAGAAAACACTCGGCTGGGAGGCCTCCACCGAGACGATGTACCCCGCTGACGGCGGC
CTGGAAGGCGCATGTGACATGGCCCTGAAGCTCGTGGGCGGGGGCCACCTGATCTGCAACCTTGA
GACCACATACAGATCCAAGAAACCCGCTACGAACCTCAAGATGCCCGGCGTCTACAACGTGGACCA
CAGACTGGAAAGAATCAAGGAGGCCGACGATGAGACCTACGTCGAGCAGCACGAGGTGGCTGTGG
CCAGATACTCTACTGGTGGCGCTGGTGATGGAGGTAAATGAATTGCCCGCCAACCGCCGAGCATCG
TCCCCCC
